# Massively Parallel Binding Assay (MPBA) reveals limited transcription factor binding cooperativity, challenging models of specificity

**DOI:** 10.1101/2024.06.26.600749

**Authors:** Tamar Jana Lang, Sagie Brodsky, Wajd Manadre, Matan Vidavski, Gili Valinsky, Vladimir Mindel, Guy Ilan, Miri Carmi, Naama Barkai

## Abstract

DNA binding domains (DBDs) within transcription factors (TFs) recognize short sequence motifs that are highly abundant in genomes. *In vivo*, TFs bind only a small subset of motif occurrences, which is often attributed to the cooperative binding of interacting TFs at proximal motifs. However, large-scale testing of this model is still lacking. Here, we describe a novel method allowing parallel measurement of TF binding to thousands of designed sequences within yeast cells and apply it to quantify the binding of dozens of TFs to libraries of regulatory regions containing clusters of binding motifs, systematically mutating all motif combinations. With few exceptions, TF occupancies were well explained by independent binding to individual motifs, with motif cooperation being of only limited effects. Our results challenge the general role of motif combinatorics in directing TF genomic binding and open new avenues for exploring the basis of protein-DNA interactions within cells.

## Introduction

Transcription factors (TFs) regulate gene expression by binding to short DNA motifs in gene regulatory regions. In bacteria, TF binding motifs are sufficiently long to provide precise addressing within the genome (1). However, in eukaryotes, binding motifs are shorter while genomes are longer, resulting in a vast number of motif occurrences, only a fraction of which are bound(1–4). TF binding in eukaryotes, therefore, depends on DNA properties that distinguish bound motif sites from unbound ones (3, 5–7).

It is commonly assumed that TF-bound motifs are distinguished by their proximity to other nearby motifs(3, 7, 8). This model postulates that DNA binding is stabilized by co-binding with other TFs at sites containing the respective motif combinations. Indeed, motif combinations are less likely to occur randomly, thereby increasing the precision of genome addressing. This combinatorial TF binding model is supported by the abundance of closely spaced motifs within regulatory enhancers(9), which is indeed required for motif-based binding cooperativity, although it could also serve other purposes, including sensing multiple TF-transduced signals and post-binding recruitment of cofactors(10, 11)

TF combinatorial binding could result from direct interactions, as exemplified by obligatory homo- or heterodimer TFs, although these typically bind at composite motifs that are still short and of low information. It is also notable that while composite motifs are characterized by a fixed distance between their half-sites, the spacing, and orientations of nearby motifs found within regulatory regions diverge rapidly(4, 9, 10, 12–16). These observations motivated an alternative model, suggesting that TFs interact indirectly by evicting the surrounding nucleosomes or by changing the DNA shape(17). The potential for nucleosome-based cooperation was demonstrated using an *in-vitro* reconstituted system with synthetic sequences(18), but its broad application in genomes has not yet been tested.

The compact and well-annotated budding yeast genome offers a convenient platform for testing the role of motif combinatorics. To this end, our lab has generated a compendium of binding data for 141 (95%) budding yeast TFs, all measured under the same conditions(19–21), which now allows us to define regulatory regions bound by multiple such TFs within cells.

Analyzing motif interactions within regulatory regions requires comparing the *in vivo* binding of a specific TF to a library of DNA variants, systematically perturbing all motif combinations (Figure 1A). Here, we introduce Massively Parallel Binding Assay (MPBA), a sequencing-based method that enables measuring TF binding to thousands of designed DNA sequences within cells using a rapid, temporally controlled readout. We applied MPBA to measure motif interactions within dozens of 164 bp-long regulatory regions, each containing a total of 3-7 motifs bound by multiple TFs. We describe some cases of cooperative interactions but find those to be the exceptions. Rather, TF binding was explained primarily by the independent contributions of individual, mostly self-preferred, motifs. We discuss the implications of our results for understanding how TFs detect their binding sites within genomes.

**Figure 1.**
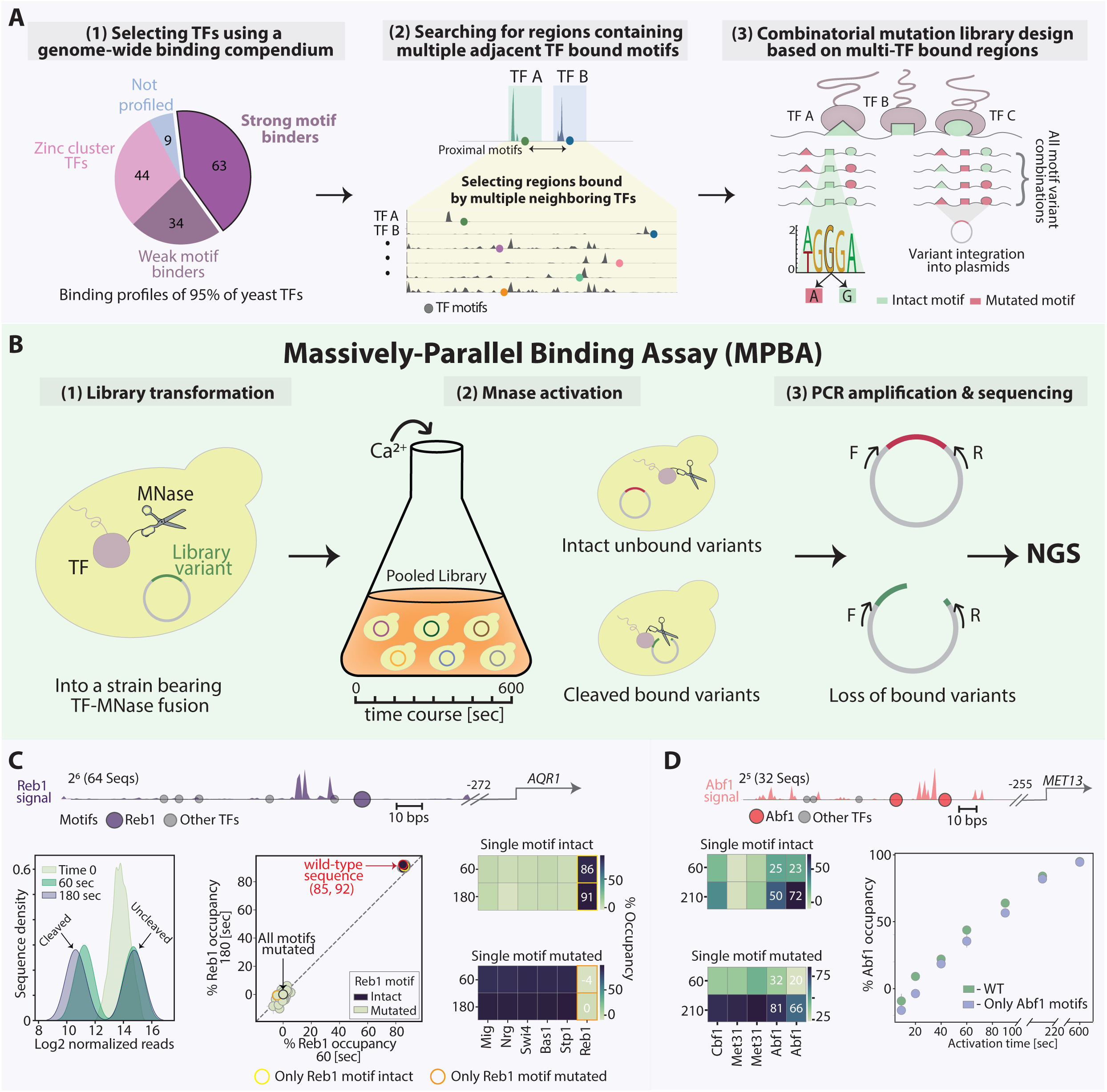
MPBA allows the sensitive detection of TF binding to a library of DNA sequence variants within cells. *A. Selecting TFs and regulatory regions for testing motif interactions:* To select candidates for cooperation, we searched for regulatory regions that contain multiple motifs bound by different TFs. For this, we used our lab dataset, which describes the genome-wide binding profiles of 145 (95%) of budding yeast TFs, collected using ChEC-seq. From those profiles, we focused on the 63 TFs showing a strong binding preference for their *in-vitro* motif (1), leaving out weak motif binders and the fungi-specific zinc-cluster TFs localizing to low-information motifs. For each of these TFs, we then selected the top-bound promoters and filtered them by the presence of the respective TF *in vitro* preferred motif showing increased signal at its proximity. We screened those regions manually to detect 164 bp segments that include not only the motif bound by the TF that directed us to this region, but also motifs associated with other TFs, showing enriched signal in their respective binding profiles (2). Each of the selected regulatory regions was used to generate a library of sequence variants, which includes the wild-type sequence and all combinations of mutated motifs, as demonstrated (3). For this, we defined for each motif the central position with the highest information content based on the TF *in-vitro* probability weight matrix (PWM; Methods). For this position we considered two variants: Leaving it intact or generating a mutation, the latter expected to abolish the binding of the associated TF. *B. Experimental procedure of MPBA:* To measure the *in-vivo* binding of a TF of interest to a library of DNA sequence variants we followed the steps shown. First, we fuse the TF of interest to a Micrococcal Nuclease (MNase). Cells containing this TF-MNase fusion are then transformed with a library of plasmids, each containing one DNA sequence variant (1). Next, cells are subjected to a short pulse of calcium which activates the MNase (2). Sequences bound by the TF-MNase fusion are then cleaved, while the unbound variants remain intact. PCR amplification is used to enrich for uncleaved sequences, whose abundance is defined using high-throughput sequencing and compared to their abundance prior to MNase activation (3). Of note, multiple libraries based on distinct regulatory regions can be transformed into the same TF-MNase bearing strain and measured in a single experiment. *C. Reb1 binding at the AQR1 regulatory region depends on its in vitro defined motif:* The *AQR1* regulatory sequence is shown on top, with all detected TF-bound motifs considered in our library design indicated in circles. The Reb1 ChEC-Seq genomic binding signal at this region is shown in purple (Methods). The distance of the selected region from the *AQR1* transcription start site (TSS) is also noted. A library containing all motif mutation combinations was transformed into a strain bearing a Reb1-MNase fusion. Shown in the lower panel are the analysis steps used to define the Reb1 occupancy at each sequence. First, we examined the distribution of sequence reads before and after triggering MNase based cleavage (left). Next, the post-cleavage abundance of each sequence was normalized by its pre-cleavage abundance (time 0) and then by the fully mutated sequence, converting normalized abundance to occupancy (middle; Methods). Shown are the Reb1 occupancy values at each of the library variants, comparing two MNase activation times (60 and 180 seconds). The intact sequence, containing all motifs, and the fully mutated sequence, lacking all motifs, are indicated by color (red and black respectively). As a further visualization of this data, we plotted the occupancy at sequences containing a single intact motif (top right; the variant in which only the Reb1 motif is intact is marked in yellow) and the occupancy at sequences in which only a single motif is mutated (right bottom; the variant in which only the Reb1 motif is mutated is marked in orange). Note that in the example shown, the single self-motif fully explained the high (>90%) Reb1 occupancy of this region. *D. Abf1 binding to the MET13 regulatory region depends on its own motif and increases over time:* Shown on top is the *MET13* regulatory region, presentation as in Figure 1C, with the Abf1 genomic signal presented in red. Shown on the left, are the Abf1 occupancies at sequences in which only one motif is intact (top) and ones in which only one motif is mutated (bottom). Shown on the right is the Abf1 occupancy at the sequence containing only its motifs intact (blue) and at the fully intact sequence (green), as a function of MNase activation time. Error bars indicate the standard error of the mean (SEM) between experimental repeats.

## Materials and methods

### MPBA

#### Library sequences and information

The information of genomic loci, the variable positions of all libraries and annotated motifs are found in Supplementary Table 1.

#### Plasmids

Plasmids used for MPBA were based on pRS316 plasmid and can be found in Supplementary Table 3. and CsiI cutting sites were added to the pRS316 using restriction free cloning and this was used as our main working plasmid.

#### Library plasmid context

For changing the surrounding library context, four genomic loci bound by Msn2 were chosen, based on their different nucleosome architecture and the total number of TFs bound. In each context, the central 120 bp promoter region containing the Msn2 binding sites was removed and replaced with SgsI and CsiI cutting sites. Next, the remaining 2 kb surrounding the binding site (1 kb upstream and 1 downstream) were introduced with point mutations to edit any existing SgsI/CsiI binding sites. In addition to these four genomic loci, another synthetic context was planned, in which each side of the cloning site has 200 bps with a low GC content (20%), aimed to evict nucleosomes. All contexts were ordered from IDT (gBlock), and integrated into the pRS316 plasmid between the XhoI and SpeI cutting sites. The context constructs can be found in Supplementary Table 3.

#### Library cloning

Libraries were ordered from IDT (4 nmole Ultramer™ DNA Oligos) and resuspended in 40 ul of Tris pH 8 (to a final concentration of 100 uM). To amplify the libraries, we first calibrated the PCR reaction (Supplementary Table 4, primers 1-2) using 4 dilutions for each library (reaching to final reaction concentrations of 44.4 nM, 4.4 nM, 1.8 nM, 0.9 nM) in order to avoid non-specific amplification products. The PCR reaction was done using Herculase II Fusion DNA Polymerase (Agilent, 600677) and was performed according to the recommendation of the manufacturer for 11 cycles. The concentration yielding the sharpest and most specific band was chosen for each library. This reaction was repeated twice more (a total of 3 reactions for each library to minimize PCR biases. reactions were combined and cleaned using the QIAQuick PCR purification kit (QIAGEN, 28104). Concentrations were measured using nanodrop machine and ranged 20-120 ng/ul.

#### Digestion

Plasmids and libraries were digested using the restriction enzymes SgsI and CsiI (Thermofisher, FD1894 and FD2114 respectively). For the restriction, we used 180 ng of the library and 2ug of our 5kb plasmid. Plasmids went through an additional step of 5’ dephosphorylation using FastAP Thermosensitive Alkaline Phosphatase (Thermofisher, EF0654) to avoid self ligation. Their cleavage was validated by comparing them to an uncut control plasmid using gel electrophoresis. Reactions were performed according to the recommendation of the manufacturer. Cleaved plasmids and oligos were then cleaned using the QIAQuick PCR purification kit, and DNA concentrations were measured (2.5-10 ng/ul for libraries and 50-100 ng/ul for plasmids).

#### Ligation

Libraries and plasmids were ligated together using the fast-link DNA digestion kit (Biosearch technologies, LK0750h) in a respective ratio of 2:1, using 225 ng plasmid, and according to the manufacturer recommendation. Ligations were cleaned using the MinElute PCR Purification Kit (QIAGEN, 20-28004), with an additional wash with the PE buffer to avoid high salt concentration. Ligated plasmids were elutied twice to increase yield.

#### Bacterial transformation

E. cloni® Electrocompetent Cells (Biosearch technologies, LC601172) were used according to the provided protocol with some modifications. Cells were thawed over ice, and the entire process (until the recovery step) was performed in the cold (4°C). After the recovery step, cells were directly inoculated in 250 ml warm LB media, and grown overnight (shaking, 37°C). A small amount of cells were plated for counting the total number of transformants (5-30M for transformants for each library). 16-20 colonies from each transformation were collected and library insertion was validated using PCR (Supplementary Table 4, primers 3-4). The integration success rate was >95%.

#### Plasmid extraction

Plasmids were extracted using the NucleoBond Xtra Maxi kit (Machery-Nagel, 740414.10), according to the provided protocol, yielding 1500 ng/ul on average.

#### Yeast transformation

To transform yeast with plasmids carrying the library variants the BY4741 strain, of genotype MATa his3-Δ1 leu2-Δ0 lys2-Δ0 met15-Δ0 ura3-Δ0, bearing a TF-MNase fusion was transformed using the LiAc/SS DNA/PEG method (22), with several modifications to increase transformant number and avoid library complexity issues. Briefly, the relevant yeast strain was freshly thawed from a frozen stock, plated on YPD plates, and grown at 30°C. A single yeast colony was inoculated in fresh liquid YPD, and grown to saturation overnight. The next morning, 250 ul of the stationary culture was diluted into 12.5 ml YPD media and grown at 30°C for 4-5 hours. The cells were washed with DDW and then with LiAc 100 mM, and resuspended in a transformation mix, containing 33% PEG-3350, 100 mM LiAc, single stranded salmon sperm DNA, 10 ug of library plasmids and 700 ng of the plasmid containing the poorly bound sequence control. Of note, while multiplexing different libraries to the same transformation, concentrations were normalized according to the complexity of each library, based on the number of variable positions it contains. The cells were incubated at 30°C for 30 minutes followed by a 30 minutes heat shock at 42°C. After the heat shock, the clles were pelleted and resuspended in 1 ml SD-URA media. 1:500 of the resuspended cells were further diluted and plated for calculating the number of transformants. These transformations yield ranged between 200K and 4M. The rest of the cells were inoculated in fresh 25 ml SD-URA and were grown to stationary phase for 72 hours (shaking at 30°C).

#### Genomic library integration

First, the HO coding sequence in the Msn2-MNase expressing strain was replaced by a short target sequence (GGTGTGGGTTTAGATGACAAGGG) using the CRISPR-Cas9 pbRA89, a gift from James Haber. Next, after testing positive using Sanger sequencing, the pbRA89 plasmid was lost from cells by growth in YPD and selection for colonies without bRA89-encoded Hygromycin resistance. Then, the libraries were PCR amplified using homologic HO locus sequences flanking the primers (Supplementary Table 4). The PCR-amplified libraries were then transformed together with the bRA89 plasmid, this time containing a gRNA directed to the short sequence introduced in the first transformation. Cells were grown in liquid YPD media containing 50 mg/ml Hygromycin to select for positive colonies containing the library embedded within the HO locus.

#### Experimental procedure

Library-transformed stationary culture was diluted into 10 mL of fresh SD-URA media (per sample) to reach OD_600_ of 4 after ~10 cell divisions at 30°C. Note, that this method obliges to compare the change of the variant pool following MNase activation. Therefore, at the beginning of the experiment, each sample was splited into two 5 ml samples that were subjected to different treatments: The non-activated sample (time point 0) will be taken directly to DNA extraction (see below), while the other sample will be subjected to MNase activation (see below), until the proteinase K digestion step, and only then will be DNA extracted.

#### MNase activation

This procedure was performed as previously described. In brief, cultures (OD600=4) were pelleted at 1500 g and resuspended in 1 mL Buffer A (15 mM Tris pH 7.5, 80 mM KCl, 0.1 mM EGTA, 0.2 mM spermine, 0.5 mM spermidine, 1 × Roche cOmplete EDTA-free mini protease inhibitors, 1 mM PMSF), and then transferred to deep well plate. Cells were washed twice more in 500 μL Buffer A, pelleted, and resuspended in 150 μL Buffer A containing 0.1% digitonin. Then, cells were transferred to a 96-well plate (PCR-96-FLT-C, Axygen) for permeabilization (30°C for 5 min). CaCl2 was added to a final concentration of 2 mM. Mnase was activated for different times, as specified in the text. Next, 100 μL of stop buffer (400 mM NaCl, 20 mM EDTA, 4 mM EGTA and 1% SDS) were mixed with 100 μL of sample. Proteinase K was then added, and incubated at 55°C for 30 min.

#### DNA extraction

Plasmid DNA was extracted both for MNase-activated cells and non-activated cells using the MasterPure Yeast DNA Purification Kit (Lucigen Corporation, MPY80200), with some modifications. Briefly, 300 ul of lysis buffer supplemented with RNase A (as recommended) were added to each sample. For MNase-activated cells, 300 ul lysis buffer were added directly to the proteinase-K treated samples. For non-activated samples, the OD600 4 was first pelleted in 1500 g to remove SD-URA media, then cells were resuspended in 300 ul lysis buffer. Following 15 minutes of incubation in 65 C, cells were moved into LoBind microcentrifuge tubes (Eppendorf®, 022431021) containing 0.5mm Zirconium Oxide beads (ZrOB05, Next Advance) and blended using the Bullet Blender 24 (Next Advance) for a single cycle of 3 minutes in level 8. Samples were cooled over ice and lysates were moved to 1.5 ml tubes, vortexed, and supplemented with 300 ul MPC buffer. Samples were vortexed vigorously and centrifuged for 10 minutes (17,000g). Supernatant was moved to 1.5 ml tubes containing 500 ul isopropanol. Samples were vortexed vigorously and DNA was precipitated (10 minutes, 17,000g). Isopropanol was removed, and pellets were washed in 70% EtOH, then air dried and finally resuspended in TE buffer.

#### Library preparation

Samples were 1.5 X SPRI (AMPure XP, A63881) cleaned and resuspended in 30 ul elution buffer (10mM Tris-HCl, pH 8) and then diluted 1:20. 1 ul was used as a template for PCR reaction for amplifying noncleaved variants and sample barcoding (Supplementary Table 4, primers F: 5-9, R: 10-41; 28 cycles) using the Phusion® High-Fidelity PCR Master Mix (NEB, M0531L). Samples were verified using gel electrophoresis, and 0.5 reversed-SPRI cleaned. DNA concentrations were measured using Qubit™ Flex Fluorometer (Invitrogen), and samples were normalized to equal amounts, then pooled together. 1 ng was taken as a template for a PCR reaction (Supplementary Table 4, primers F: 42-65, R: 66-77) using KAPA HiFi HotStart ReadyMix (Roche, KK2602) for adding IIlumina indices and machine adapters. Samples were 1X SPRI cleaned, then concentrations and library quality were validated using Qubit and tape station (Agilent).

#### Sequencing

Libraries were sequenced using the NovaSeq 6000 machine. Runs were performed with the SP200 kit (20040719), parameters: R1 - 150 cycles, Index1 - 8 cycles, Index2 - 8 cycles, R2 - 6 cycles. In each run, 5% PhiX DNA was added to increase complexity.

### ATAC-seq

#### Experimental procedure

Cells were grown as in MPBA, to reach OD_600_=4 and were then pelleted in 1.5 ml Eppendorf tubes at 1500 g for 60 seconds and washed twice with 200 μL Spheroplasting Buffer (SB) (40 mM HEPES [pH 7.5], 10mM MgCl_2_ 1M Sorbitol). The cells were next resuspended in 1 μL Lyticase 10ku (Sigma, L2524) dissolved in 199 μL SB, transferred to a 96-well plate, and incubated for 30 minutes at 30°C. The spheroplasted cells were pelleted at 1500 g for 60 seconds, washed twice with SB as in the previous step, resuspended in 2 μL of RNase-A (5 μg/μL) dissolved in 98 μL SB, and incubated for 30 minutes at 37°C. Cells were then pelleted and washed twice as in the previous steps, resuspended on ice in 25 μL Transposase reaction mix (Tn5 transposase enzyme produced in house [6.5μM] loaded Tn5MEDS-A/B oligonucleotides (Supplementary Table 4) (23), 2x Tn5 Buffer [10 mM Tris HCL PH 7.6, 10 mM MgCl_2_, DMF in 1:4 ratio], DDW) and transferred to a 96-well plate for 60 minutes incubation at 37°C. Transposase reaction clean-up mix (5% SDS, 20mg/ml Proteinase K, Clean-up buffer [NaCl 5M, EDTA 0.5M, DDW]) was added, and plate was incubated for another 30 minutes at 40°C. Samples were 2x SPRI (AMPure XP, A63881) cleaned and resuspended in 22 μL elution buffer (10mM Tris-HCl, pH 8).

#### Library preparation

The eluted samples were used as a template for a PCR reaction for amplifying Tn5 tagmented fragments and sample barcoding (2× Kappa HIFI [Roche], 5 μL nextera primers F: i5, R: i7; 12 cycles; see primer list in Supplementary Table 4). Next, samples were 0.65x reversed SPRI cleaned and resuspended in 22 μL elution buffer (10mM Tris-HCl, pH 8), then concentrations and library quality were validated using Qubit and tape station (Agilent).

#### Sequencing

Libraries were sequenced using the NovaSeq 6000 machine. Runs were performed with the S1 kit (20028319), parameters: R1 - 61 cycles, Index1 - 8 cycles, Index2 - 8 cycles, R2 - 61 cycles.

### Quantification and statistical analysis MPBA Pipeline

Our pipeline was built using SnakeMake(24) and can be found in [GitHub](https://github.com/sagieb/MPBA). Briefly, the forward library primers were removed using cutadapt(25). Then, AdapterRemoval(26) was used to demultiplex each 32-sample pool into separate samples. Next, in order to demultiplex the different libraries in each sample, each read was assigned to the right library using cutadapt. Reads with improper 3′ prime alignment were filtered out in order to remove sequences containing indels. Then, each combination of variable positions within each library was counted. In addition, the number of reads in each sample aligning to the poorly bound sequence control was counted.

### MPBA Data filtering

Variable position combinations that do not match the library were removed. Samples quality was verified by repeat similarity and read count. Samples with low sequence depth and/or low correlation to other technical repeats were removed from further analysis.

### MPBA Data normalization

The read count of each sample was normalized to 10^6^ to account for sequence depth. Next, technical repeats were averaged and log2 transformed. Then, each MNase-activated sample was normalized to the corresponding pre-activated sample. Lastly, biological repeats were averaged.

### Occupancy measure

To calculate the relative occupancy, each time-point zero normalized sequence was normalized to the fully mutated sequences and converted to % using the following equation:

(1-(0.5^-(value of tp0 normalized sequence - tp0 normalized fully mutated sequence value)^)).

### Normalized occupancy measure

To calculate histone occupancy compared to the genome, a parameter representing the highest nucleosome occupancy of library-sized promoter region (164 bps) was calculated. To this end, the MNase-seq(27) signal was summed over all library-sized regions promoter, using a sliding window of 10 bps. The top 1% of this calculated distribution served as the value of a 100% nucleosome occupancy. The ratio between the MNase-seq signal over each library to this value was calculated and served as the genomic occupancy over each tested region. For calculating the nucleosome occupancy over each library sequence, as presented in Figure 4G, the genomic occupancy and the fold change between each sequence and the wild-type one was used, as follows:

0.5^(value of sequence normlized to tp0 & wild−type variant value) ∗ (Library nucleosome occupancy)^

### Cooperativity score

For a given pair of motifs, the occupancy (in fraction) of each motif in the absence of the second motif was calculated in each context (all combinations of all non-pair motifs found in the respective library). Specifically, for each pair in each context, the log2 fold-change between the sequences containing only one intact motif to the sequence where both are mutated was calculated and converted to occupancy (as in the Occupancy measure section). Then, the expected cooperativity, assuming independence, of the motif pair was calculated for each context as follows:

1 - (1-Occupancy_motifA_) * (1-Occupancy_motifB_).

The observed motif pair occupancy was calculated for each context using the log2 fold change between the sequence in which both motifs are intact to the one where both motifs are mutated. The cooperativity score is the average difference between the observed and expected occupancy over all contexts.

### Genome-wide promoter binding

ChEC-seq Samples were processed as in (21). Promoters were defined as in (19). Each promoter was assigned with the average sum of signal it received over all repeats.

### Library region genomic binding

For the correlation heatmap presented in Figure 6A, the sum of signal received for each TF over each genomic region used to generate libraries was summed. Next, the sum of signal received for all these library regions was correlated between the different TFs. For the region binding strength heatmap presented in Figure 6B, the signal received over each genomic region representing a library was summed and divided by the highest signal received on a similar-sized region (164 bp) over the entire genome of the respective TF.

### Motif enrichment

For the motif enrichment analysis, all possible 7-mer sequences were given a numerical index (Forward and reverse complement forms of each 7-mer were given the same index). Each nucleotide in the yeast genome was indexed according to the 7-mer that begins from it. To score each 7-mer occurrence, the signal around its mid-position was averaged (20 bp window). The averaged signal for each 7-mer was then calculated across all of its occurrences in all promoters and was assigned as its relative binding score. Next, the relative binding score of all 7-mers containing the relevant TF motif (Ste12 - TGAAAC, Tec1 - GAATG[CT], Reb1 - CGGGTAA, Supplementary Table 2) was averaged and divided by the average signal of all 7mer combinations.

### Motif Euclidean distance

The Euclidean distances between the motifs of the TFs presented in Figure 6G were calculated using the Probability weight matrices (PWMs)(28) from same as in (3).

### Non-cooperative recruitment

To measure the non-cooperative recruitment presented in Figures 5A and 6F, the average effect of each non-canonical motif in each library was calculated. In detail, after data normalization, the average log2 fold-change of all sequences in which the respective non-canonical motif is intact and the self-TF positions are mutated was normalized by the average log2 fold-change of all sequences in which the respective non-canonical motif is mutated and the self-TF positions are mutated. Then, the relative position occupancy was calculated as in the Occupancy measure section.

### Self-motif-dependent region occupancy

To calculate the self-motif-dependent region occupancy presented in Figure 6E, after data normalization, the average log2 fold-change of all sequences in which all self-TF motifs are intact was normalized to the log2 fold-change of all sequences in which all self-TF motifs are mutated. Then, the relative occupancy was calculated as in the Occupancy measure section.

Number of TFs bound to each context:

For this aim, we used our lab data containing the genomic binding signal of most yeast TFs. For each TF, the sum of normalized signal received on each promoter was calculated and z-score transformed. This data is found in Supplementary Table 5. Then, the top number of TFs bound to the central promoter found in each context was calculated, asking how many TFs received a z-score >3.

### ATAC seq

Reads were aligned to the *S. cerevisiae* genome (SacerR64) using bowtie2(29, 30) with parameters: --very-sensitive --trim-to 30 –dovetail. Reads larger than 140 bps were discarded using genome coverage(31). The coverage was calculated on the 5’ position of each read. The repetitive region in chromosome 12 (451600:489500) was removed and the reads of each sample were normalized to 10^7^.The presented data was smoothed using 20 bps window size in all figures, except from Figure 4F in which 50 bps window was used.

### In-vitro and In-vivo MNase-seq

The “InVitro” and “YPD” normalized data samples were taken from(32). The genomic annotations were converted to the SacerR64 genome using CrossMap(33).

## Results

### Massively Parallel Binding Assay (MPBA) for rapid screening of thousands of DNA variants

We developed MPBA for comparing the binding of a given TF to a library of designed DNA sequences. This method follows the conceptual design of Massively parallel reporter assays (MPRA)(34–38), but replaces the expression-based output of MPRA by a more direct readout of TF binding to DNA.

In MPBA, the endogenous TF of interest is fused to a Micrococcal Nuclease (MNase), which cleaves TF-bound DNA upon a short calcium pulse, as described in previous specific applications (39). We adapted this method to enable variant screening as follows (Figure 1B). First, we integrate our library variants into a specialized plasmid and transform the plasmid into cells carrying the TF-MNase fusion. Next, we activate the MNase to trigger the cleavage of TF-bound plasmids. Following this step, the library variants are PCR amplified, thereby capturing only the non-cleaved (unbound) sequences. Then, the frequency of each variant is defined using high-throughput sequencing. Finally, we normalize the abundance of each sequence to its abundance prior to MNase activation and then to that of the fully mutated sequence, lacking all motifs. This provides a measure for TF occupancy at each sequence within the library, namely the proportion of cells in which this sequence is bound.

### MPBA captures motif binding and nucleosome eviction by the General Regulatory Factors

Plasmids provide a convenient platform for massive parallel analysis of sequence libraries but might be limiting for capturing properties of TF binding inside cells, given their uncertain chromatin environment. Budding yeast differs from higher eukaryotes in lacking large-scale closed chromatin domains. Still, nucleosomes might be more loosely packed on a plasmid, which would reduce cooperative effects, or, conversely, might be denser, limiting TF binding. As a first test of our method, we therefore tested the nucleosome-organizing general regulatory factors (GRFs) Reb1, Rap1, and Abf1 (40, 41), asking whether MPBA can capture their tight motif binding and their role in nucleosome eviction. Of note, when compared to other TFs, the GRFs bind a large fraction of their motif sites, the majority of which are present in promoters deprived of other TFs (Figure S1A).

We selected six 164 bp GRF-bound genomic loci, biasing towards the limited fraction of regions bound by additional TFs (Figure S1B). Six respective sequence libraries were then designed as follows. First, for each of the *n* TF-bound motifs present in the selected region, we considered two variants: the native motif and a 1-bp motif mutation abolishing TF binding (Figure 1A; Supplementary Tables 1-2). Each library then includes all combinations of these intact or mutated motifs, totaling 2^n^ sequences. For the GRFs, the six libraries include 32-128 sequences corresponding to the 5-7 TF motifs present in the selected regions (Figure S1B).

The designed libraries were integrated into a specialized plasmid (Supplementary Table 3), multiplexed, and transformed into cells carrying the respective GRF fused to an MNase (Figure 1B). As expected, MNase activation led to rapid and reproducible depletion of sequences containing intact GRF motifs, while sequences lacking the motif remained at high abundances (Figures 1C and S1C). In the case of Reb1, the depletion at 60 seconds of MNase activation translated into estimated occupancies of 85-92% at the wild-type sequence containing all intact motifs, which increased to a maximum of 95% in three minutes. The binding of Rap1 and Abf1 was again dependent on their known motifs, reaching somewhat lower estimated occupancies at similar activation times (3 minutes, 40-80% at the fully intact sequence). To verify that this reduced depletion does not reflect a limitation of our system, we performed a tight time course measuring the Abf1 binding to its respective libraries, revealing a gradual increase in Abf1 binding with the MNase activation time, reaching an almost full occupancy at the *MET13* library (100%; Figures 1D and S1C). We conclude that MPBA captures the strong GRF motif binding and further provides, for the first time, a time-dependent measure of the fraction cells in which this motif is bound.

We next asked whether our plasmid-based approach can also capture the GRF role in nucleosome eviction. To quantify nucleosome occupancy, we measured the binding of the histone H3 to our six libraries using MPBA (Figure 2A). As expected, in five of these libraries, H3 eviction increased continuously with increasing GRF binding (Figures 2B-C). In these libraries, the GRF-inflicted ~15-55% decrease in H3 found in our assay is compatible with the average GRF effects in the genome, as estimated by comparing *in vivo* and *in vitro* nucleosome occupancy at the GRF binding sites(32) (Figures 2D-E). We conclude that MPBA captures not only the high GRF motif occupancy but also their role in nucleosome eviction.

**Figure 2.**
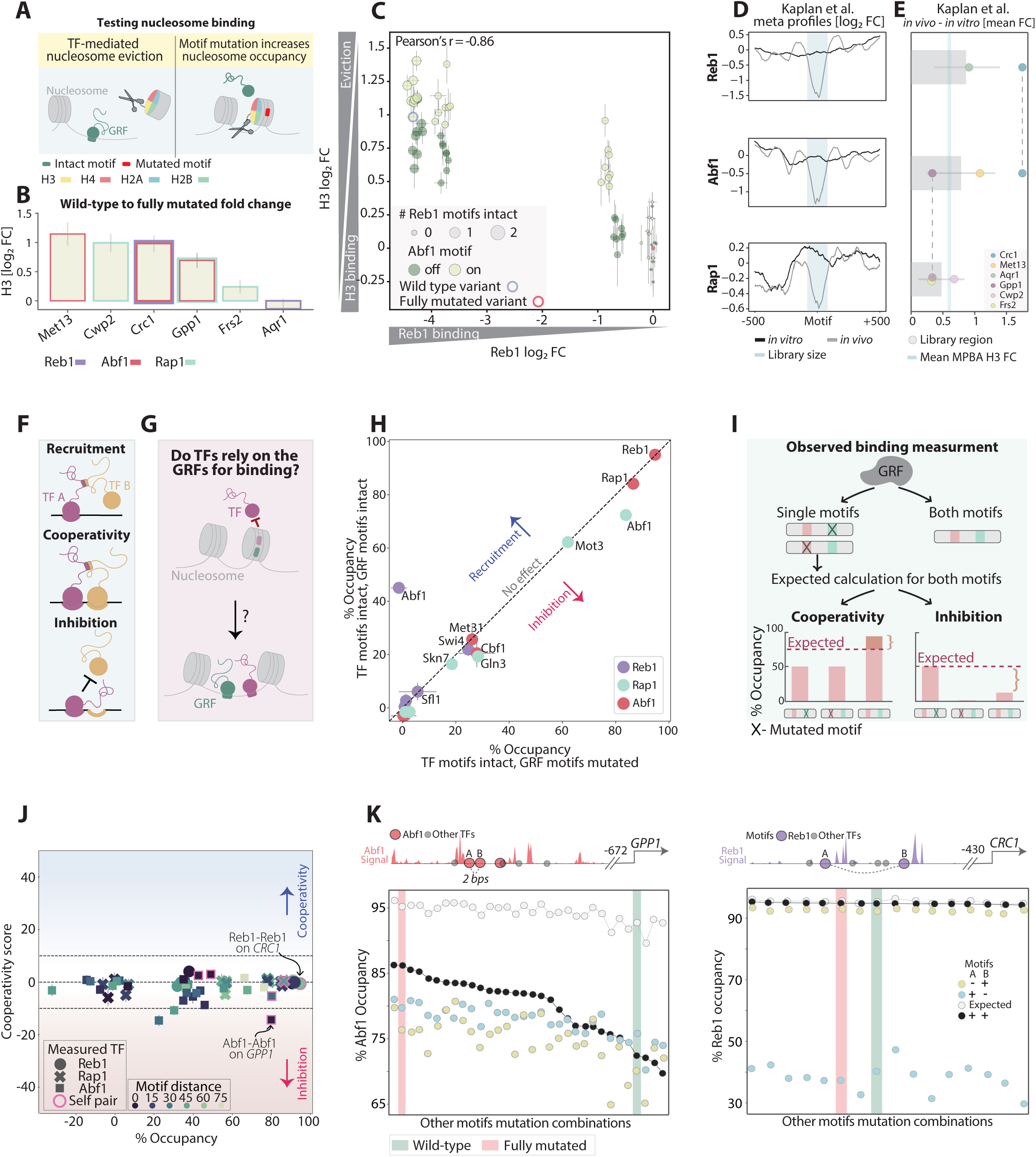
The GRFs rarely cooperate to localize to their binding sites. *A. TF mediated nucleosome eviction:* A scheme describing the scenario of nucleosome eviction as a consequence of TF binding at its binding site. H3 histone was fused to an MNase and assayed using MPBA. *B. H3 binding is increased when eliminating GRF binding:* For each GRF library, shown is the fold change of H3 occupancy between the wild-type sequence and the sequence in which all variable positions are mutated. Error bars indicating SEM calculated for all repeats (3) in each experiment. The relevant GRFs are indicated by color for each library. *C. The binding motifs of the GRFs Reb1 and Abf1 inhibit H3 histone binding to the Crc1 promoter region:* The binding at all individual sequences in the *Crc1* library is shown for Reb1 (x-axis) and H3 (y-axis). The mode (intact/mutated) of the Abf1 motif found in this region is shown in color, and the number of Reb1 intact motifs is indicated by dot size. Error bars represent the SEM calculated for all repeats. The wild-type and fully mutated sequences are marked in blue and red, respectively. Note the inversed correlation between Reb1 and H3 binding, reflecting the Reb1 function in nucleosome eviction and the additional nucleosome depleting effect attributed to Abf1 binding (color). *D-E. The effect of the GRFs on nucleosome binding corresponds to the difference between in vivo and in vitro nucleosome occupancy:* For each GRF, shown in (D) is the mean *in vivo* and *in vitro* signal around the 200 most bound motif occurrences in promoters, measured by Kaplan et al (32) (motifs were defined in Supplementary Table 2; see Methods for promoter definition). The mean fold change between these signals is summarized in (E). The fold change for each tested library is shown in dots, and the mean fold change over the libraries relevant to each GRF is indicated in red. The blue line indicates the mean nucleosome binding fold change between the wild type and the fully mutated sequence for all MPBA-GRF tested libraries. *F. Cross-motif dependencies:* A scheme demonstrating 3 different modes of TF interactions. While recruitment (top) can occur independently of proximal motifs, cooperativity (middle) and inhibition (bottom) require an interaction between nearby binding sites. Only the two latter modes are able to increase TF genomic binding specificity. *G-H. The Binding of the GRFs has little effect on TF binding to proximal motifs:* TFs may rely on the GRFs for binding at their own motifs as the GRFs evict nucleosomes (B). To examine this, we measured the binding of 10 additional TFs whose motifs are present within the designed libraries shown in Figure S1B. (C) To measure the effect of the GRFs on the binding of those other TFs (or of other GRFs), we compared the mean TF occupancies (at 180 sec MNase activation time) at sequences containing (y-axis), or lacking (x-axis) the respective GRF motifs, indicated by color. These comparisons reveal only a single case of recruitment, in which a Reb1 motif allows Abf1 occupancy at the *CRC1* regulatory region, and some cases of mild inhibition, including the inhibitory effect of the Rap1 motifs found in the *GPP1* regulatory region on the Abf1 binding. Overall, the binding of all measured TFs appears to be independent of the GRF binding. *I. Defining motif-cooperativity score:* By motif cooperation, we refer to cases in which the presence of a motif pair stabilizes the binding of a TF, beyond the individual contributions of each motif. To measure cooperation, we first defined the expected occupancy for sequences containing a pair of motifs, using sequences with only one of the two motifs intact, assuming independent binding to each individual motif (Methods). In the schematic example shown on the bottom left, sequences in which only one of the motifs is present while the other is mutated reach 50% occupancy. Accordingly, the expected occupancy of sequences in which both motifs are intact, assuming independence, is 75% (dashed line; Methods). The measured occupancy of such sequences can exceed this expectation, suggesting that the motifs cooperate to stabilize the TF binding. Alternatively, the observed binding to the sequences containing both motifs can be lower than expected, indicating inhibition (bottom right). The cooperation score is defined as the difference between the measured and expected occupancy, with positive values corresponding to cooperative enhancement of binding while negative ones describe cases of inhibition (See Methods for mathematical definition). *J. GRFs binding is independent of motif cooperation:* Shown are the averaged cooperation scores as a function of the mean observed occupancy over all sequence variants containing the two individual motifs intact. Plotted are all 63 motif pairs within the six GRF libraries that contain at least one GRF motif, measured for the respective GRF. Colors indicate the distance between the motifs, where distance 0 meaning that the second motif starts one base-pair after the end of the first motif. The different shapes represent the different GRFs and self-motif pairs are marked in red. Error bars indicate the SEM (Methods). *K. Proximal Abf1 motifs demonstrate inhibitory relations:* Our analysis in D demonstrates inhibition of Abf1 binding when its two proximal motifs found on the *GPP1* promoter are intact (left). This effective inhibitory relation is visualized by examining the Abf1 occupancy of individual sequences. As shown on the library scheme (top left, presentation is as in Figure 1C), the *GPP1* library contains a total of 7 motifs. Therefore, the interaction between the two Abf1 motifs can be measured within 32 different adjacent motif combinations (x-axis). Within each such motif combination, we compared the occupancy of three mutation combinations in the two Abf1 motifs: Two that contain only one non-mutated Abf1 motif (+/− and −/+; blue and yellow) and one in which both motifs are intact (++, black). Shown on the bottom left are the occupancies of each of those three sequences within each context (y-axis), sorted by the observed occupancy when both motifs are intact. The respective expected occupancy of both motifs, assuming independence, is also shown (white). Fully intact and fully mutated motif combinations are marked by color (green and pink respectively). For comparison, the respective plot is also shown for the two Reb1 motifs within the *CRC1* regulatory region, measured with respect to Reb1 binding, which shows no cooperation (right).

### Cross-motif dependencies: Recruitment vs. Cooperation

Within regulatory sequences, TFs mostly bind to their own motif but may also depend on motifs of other TFs. In our data, this was exemplified by the binding of Abf1 to the *CRC1* regulatory region, which was mostly explained by a Reb1 motif rather than its own (Figure S1C). Given our motivation of defining the role of motif interactions in guiding TF binding preferences, we were mainly interested in such cross-motif dependencies, distinguishing between two modes: recruitment and cooperativity (Figure 2F). In recruitment, a motif that is not directly bound by the tested TF still promotes binding but independently of other motifs. This would be the case, for example, when the tested TF is recruited by a second TF that directly binds at this recruiting motif. Those cases, while interesting, do not increase specificity as TF binding still depends on a single motif. Cooperative binding, by contrast, refers to cases where two proximal motifs stabilize binding beyond their individual contributions. In these cases, specificity increases, as two nearby motifs are significantly less likely to occur at random compared to one.

The GRFs bind a large fraction of their motifs, which may indicate independent motif binding. Through their role in organizing nucleosomes, however, they may promote the binding of other TFs (Figure 2G). We examined for such cases by testing the binding of the ten additional TFs localizing to the six GRF-bound regulatory regions using MPBA. These TFs reached occupancies that were significantly lower than those of the GRFs, and in none of those cases was TF binding significantly influenced by the presence of the GRF motif (Figure 2H). In fact, the only exception was the dependence of Abf1 on the Reb1 motif at the *CRC1* promoter, noted above.

We next defined a rigorous measure of motif cooperation, which is assigned for each pair of motifs relative to the binding of a specific TF. For this, we considered the sequences in which one motif in the respective pair is intact and the second is mutated. Assuming that the contribution of those motifs to TF binding is independent, the occupancy at sequences in which both motifs are intact can be predicted. The contribution of motif cooperation is, therefore, the difference between the measured occupancy when both motifs are intact and the predicted one (Figure 2I). We note that this measure may depend on the state of other TF motifs (wild type or mutated) found within the tested region. Therefore, we average the cooperativity scores assigned to a given motif pair over all possible states (wild type or mutated) of the other TF motifs found in the library.

As expected, in the case of the GRFs, cooperation scores were low and contributed less than 10% to the total occupancy (Figure 2J). The only exception from linearity was an apparent repression, whereby the binding of Abf1 to its two adjacent motifs at the *GPP1* promoter was lower than expected (Figures 2J-K).

### Motif cooperation explains the binding of the interacting Ste12-Tec1 TFs

To verify that our method can detect binding cooperativity, we considered the two known interacting TFs, Ste12 and Tec1. The heterodimers Ste12-Tec1 bind to filamentous-related genes, while Ste12-Ste12 homodimers bind to mating-related ones (42, 43). Accordingly, Ste12 and Tec1 share common targets and, when tested across the genome, preferentially localize to the motifs of one another (Figures 3A-B). We selected seven regulatory regions bound by either or both TFs, defined the TF-bound motifs in those regions, and generated corresponding libraries containing all combinations of motif perturbations, as above (Figure 3C). Measuring Tec1 and Ste12 across those libraries revealed high occupancies at the intact sequences, reaching ~80% at the strongly bound regions, including the *PCL2* and *TEC1* promoters bound by both TFs, and the *GPA1* and *KAR4* regions bound only by Ste12 (Figure S2). Regions showing lower binding in the genome (*POP3*, *SIM1*, and *CDC6*) showed correspondingly low occupancy in our system (Figures 3A and S2).

**Figure 3.**
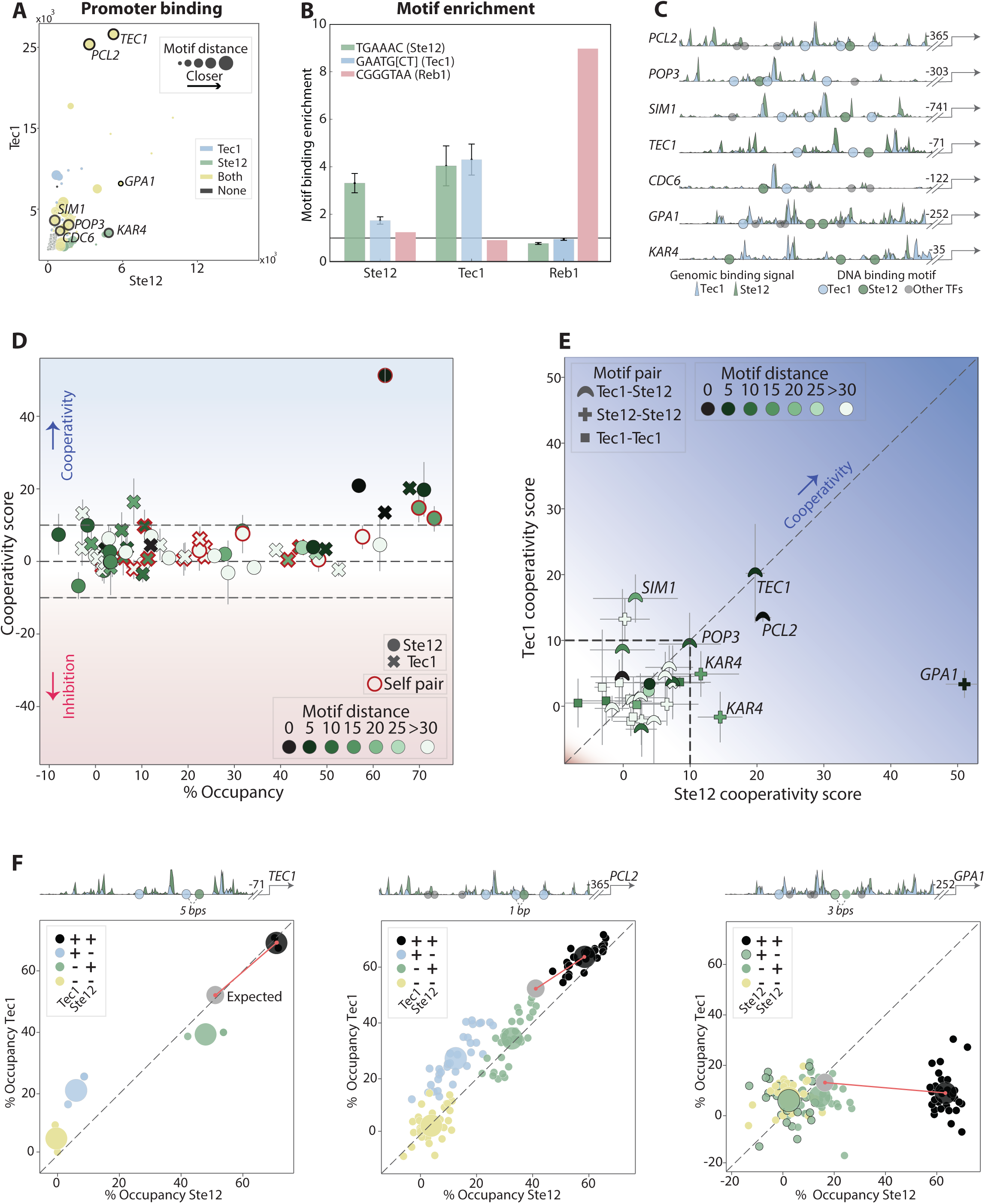
Cooperating motifs guide the binding of the interacting Tec1 and Ste12 TFs. *A-C. Selecting regulatory regions for testing motif-dependent binding of Tec1 and Ste12:* To select promoter regions bound by Ste12 or Tec1 we examined their respective genome-wide binding profiles. The scatter plot (A) compares the Tec1 and Ste12 genomic binding signal at each individual promoter, highlighting in colors which motif combination is present in the promoter and the (closest) distances between each motif pair shown by size. The selected regions are found within the promoters marked in black. The *TEC1* and *PCL2* promoters, for example, contain both the Ste12 and Tec1 motifs at close distance, while *KAR4* contains two proximal Ste12 motifs but no Tec1 motifs. When tested across the entire genome, binding locations of Ste12 and Tec1 are enriched not only in their own motif, but also in the motif of their partner, as shown in (B; Methods), with Reb1 serving as a control. The selected regulatory regions are shown in (C), indicating all motif locations mutated in our libraries and showing the genomic binding signal of both TFs in these regions. The signal of each TF is normalized to the maximum value received within each region. *D-F. High occupancy by Tec1 and Ste12 depends on cooperating motifs:* Shown in (D) are the cooperation scores, as a function of the observed occupancy when both motifs are present (presentation as in Figure 2E). Included here are all pairs within the seven libraries covering all possible Tec1 and Ste12 motif combinations. A direct comparison of the cooperativity scores of Tec1 and Ste12 received for each motif pair reveals similarity in the dependent binding to the composite Ste12-Tec1 motif (E). The pattern of motif pair cooperativity can be appreciated from the occupancy at each library sequence variant, as shown for the three indicated examples (F), with sequences colored by the mutation status of the two cooperating motifs, as indicated.

We measured cooperation scores for the 35 motif pairs covering all Ste12 and Tec1 motif combinations. Cases of strong cooperation were readily identified (Figures 3D-E). For example, the regulatory regions of *PCL2* and *TEC1* were strongly bound by both TFs, but only when both adjacent Ste12 and Tec1 motifs were present, while mutating either motif led to a strong reduction in binding (Figure 3F). Similarly, the high Ste12 occupancy at its unique *KAR4* and *GPA1* regions entailed two adjacent Ste12 motifs (Figures 3D-F). Therefore, the co-binding of the interacting Ste12-Tec1 TFs is mirrored in their motif cooperation, which is readily detected by MPBA.

### Msn2 and Sok2 occupancy at regulatory regions depends on multiple independent motifs with no apparent motif cooperativity

Motif cooperation is most likely to occur in regulatory enhancers that contain multiple TF binding motifs. Examining our lab compendium of 141 (>95%) TF binding profiles highlighted two groups of TFs that bind overlapping promoters, broadly corresponding to activators and repressors of stress genes. As representatives of those groups, we selected Msn2 and Sok2, which appeared as promising candidates for participating in cooperative binding, given that both are guided to their genomic locations through (largely disordered) regions located outside their DBDs (Figures 4A and S3A).

**Figure 4.**
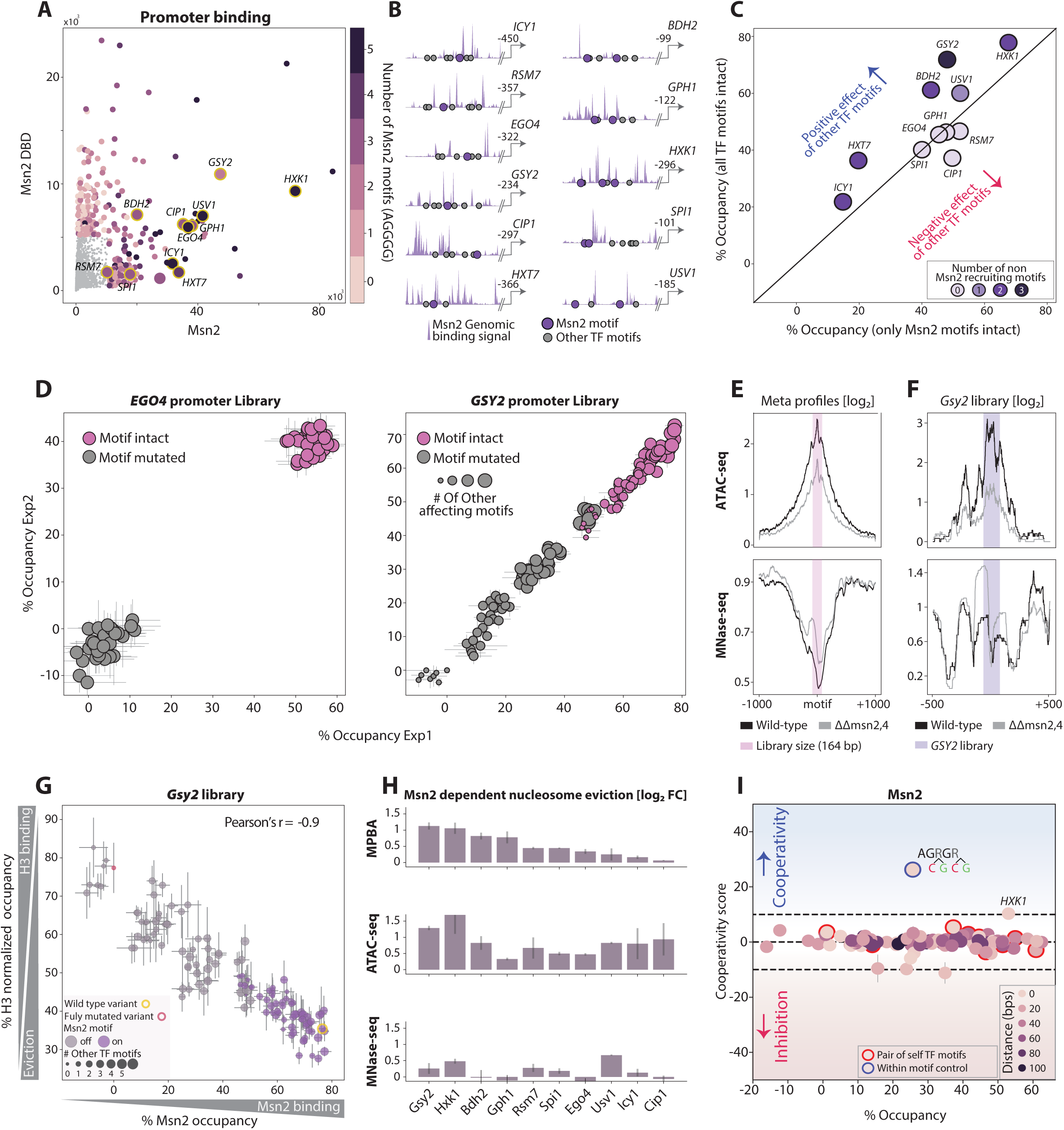
Msn2 binding is explained by additive contributions of multiple, mostly self-preferred, proximal motifs: *A-B. Selecting regulatory regions for testing motif-dependent binding of Msn2 and Sok2:* As candidates for motif-cooperation, we selected regulatory regions whose binding by Msn2 or Sok2 depends on amino-acid sequences found outside their DBDs. For this, we used our genome-wide binding dataset, in which we profiled the binding of the wild-type TFs as well as TF mutants containing only their DBDs. Shown in (A) is a scatter plot comparing the binding signal of Msn2 and its DBD-only mutant, as measured across each of the budding yeast promoters, with the number of Msn2-motifs (AGGGG) indicated by color. Regions found within the yellow-circled promoters were selected for further analysis and are shown in (B; presentation as in Figure 1C). A similar analysis and the libraries chosen for Sok2 is shown in Figures S3A-B. *C-D. Msn2 binding reaches high occupancy and is guided by multiple, mostly canonical motifs:* Shown in (C) is the occupancy of Msn2 at each intact regulatory region (relative to the fully mutated sequence), as a function of the occupancy of the same region mutated in all motifs other than the canonical Msn2 binding sites (AGGGG). Occupancy at all individual sequences is also shown for two libraries (D), with the mode (intact/mutated) of the single Msn2 motif found in these regions shown in color, and the number of non-canonical motifs contributing to occupancy indicated in dot-size (Methods). Error bars represent the SEM calculated for all repeats in each experiment (3-4 repeats for each experiment). See a similar analysis of Sok2 in Figures S3C-D. *E-H. Msn2 binding triggers nucleosome evection around its genomic binding sites:* Shown in (E) is the mean ATAC-seq and MNase-seq(27) signals at the surrounding of the 200 most bound Msn2 motifs in all annotated promoters (Methods), in wild-type cells (black) and cells deleted of Msn2 and its close paralog Msn4 (gray). The signals around the selected *GSY2* regulatory region are shown in (F). Note the increase in nucleosome occupancy (MNase-seq) and the reduction in DNA accessibility (ATAC-seq) following the Msn2/4 double deletion. The occupancy at all individual sequences in the *GSY2* library is shown in (G) for Msn2 (x-axis) and H3 (y-axis, see Figure S4A for other libraries). Msn2 binding is presented in % occupancy, as described above, while H3 binding was normalized to the genomic nucleosome occupancy (Methods). Colors indicate the mode of the Msn2 motif (intact/mutated) in each sequence, and dot sizes indicate the number of other intact TF motifs. Presented in (H) is a comparison of the Msn2-dependent nucleosome eviction around the tested regulatory regions in MPBA and in the genome (ATAC-seq and MNase-seq). For MPBA, shown is the fold change between the mean binding at sequences containing intact Msn2 effective positions (Occupancy > 10%; Methods) to the mean binding at sequences in which these positions are mutated. For ATAC-seq and MNAse-seq, the fold change between the signals in wild-type cells and those deleted of Msn2/4 is shown. *I. The binding of Msn2 is independent of motif cooperation:* Motif cooperation scores are shown as a function of the observed occupancy for all 97 motif pairs containing at least one Msn2 canonical motif within the Msn2-testing libraries following 180 seconds of MNAse activation. Color code indicates the distance between the tested motifs of each pair. The blue-circled dot is a control that includes two variable positions found within the same Msn2 motif.

We selected 18 regions, each bound by Msn2 or Sok2 and multiple additional TFs, designed respective libraries mutating all motif combinations, and applied MPBA to measure library binding by Msn2 or Sok2 (Figures 4B and S3B). In all libraries, the intact sequence was depleted, indicating ~40-80% and up to ~35% occupancies of Msn2 and Sok2 at their respective intact regions (Figures 4C and S3C). Those occupancies were mostly explained by the known Msn2 or Sok2 motifs, as exemplified by *EGO4* (Msn2) and *OCA5* (Sok2) regions (Figures 4D and S3D). In some cases, motifs that are better aligned with other TFs contributed to the occupancy, as was particularly notable in three of the tested Msn2-bound regions (*GSY2, BDH2,* and *HXT7*; Figures 4C-D).

Msn2 binding to the genome was reported to reposition or evict surrounding nucleosomes(1, 27). To test this in our assay, we applied MPBA to measure the binding of H3 across our Msn2 libraries and further quantified nucleosome occupancy across the genome in the presence or absence of Msn2 and its close paralog Msn4 by applying ATAC-seq and analyzing existing MNase-seq data in cells growing under our tested conditions(27) (OD_600_=4, Figure 4E-F). In all ten tested libraries, the H3 occupancy increased continuously with the decreasing Msn2 binding (Figure 4F-G and Figure S4, Methods). Notably, this Msn2-triggered nucleosome eviction, measured on our plasmid, was comparable to the eviction seen in the genome (Figure 4H). We conclude that MPBA simulates not only the binding of Msn2 to the tested regulatory regions but also its role in the organization of nucleosomes.

Next, we searched for cooperating motif pairs, considering those that contained at least one TF motif (97 for Msn2 and 67 for Sok2). Contrasting our expectations, Msn2 and Sok2 occupancies were well explained by independent binding at individual motifs (Figures 4I and Figure S5A). The only exceptions were the *HXK1* region, where motif cooperation increased somewhat an already strong Msn2 binding (Figures 4I and S5B), and a control sequence in which two variable positions were inserted within the same Msn2 motif, giving the expected high cooperating score (Figure 4I). To control for MNase activation, we repeated the analysis of Msn2 binding using a tight time course. However, despite the continuous increase in occupancies, cooperativity scores remained low (Figure S5C). Therefore, Msn2 and Sok2 binding at the selected regulatory sites appears independent of motif cooperation.

### Msn2 retains independent motif binding across different plasmid and genomic contexts

Msn2 binding results in significant eviction of surrounding nucleosomes but appears independent of this eviction, as indicated by the lack of significant motif binding cooperation. Still, cooperative binding that propagates through chromatin might be sensitive to the broader chromatin context extending beyond the immediate motif-surrounding sequence. Since our plasmid provides an uncertain chromatin landscape that could differ from that found in the genome, we asked whether we could retrieve motif cooperativity by changing the specific plasmid context within which we embedded our 164 bp tested regulatory regions (Figure 5A).

**Figure 5.**
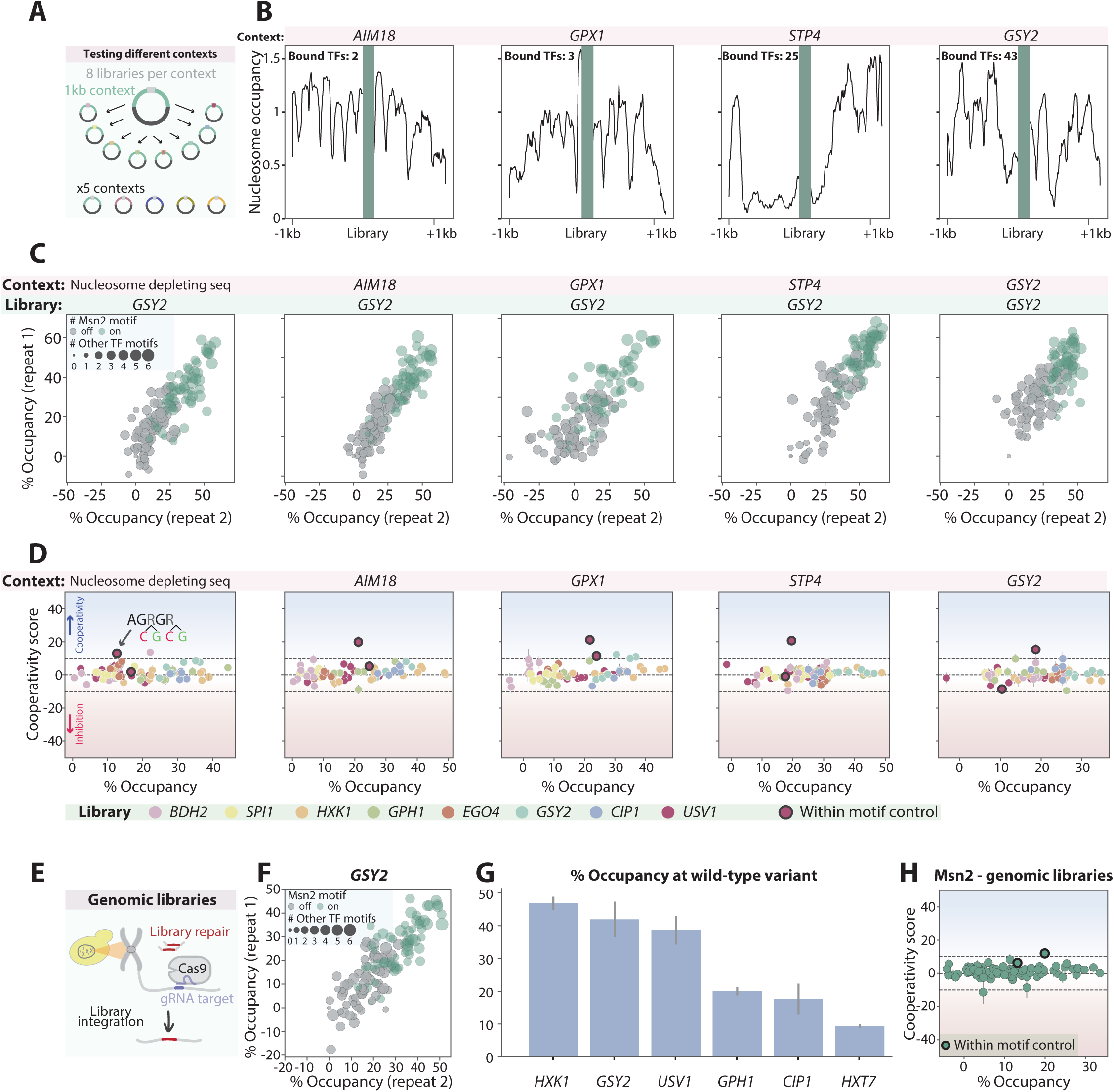
Msn2 shows no cooperativity across multiple DNA contexts: *A-B. Testing Msn2 libraries in different plasmid contexts:* A scheme illustrating the embedding of the Msn2 libraries within 5 different plasmids, each containing a distinct 2kb context. Four of the plasmids contain genomic contexts that differ in their chromatin landscape, as shown by the MNase-seq profile presented in (B), and the fifth contains a synthetic context that is expected to be depleted of nucleosomes (Methods). The number of TFs bound to the central promoter is mentioned for each context. *C. Msn2 binding to the GSY2 promoter region depends on its binding motif in all measured contexts:* Two repeats showing the binding of Msn2 to each sequence in the *GSY2* library are presented for each context. The mode of the Msn2 motif (intact/mutated) and the number of other TF intact motifs are indicated in color and dot size, respectively. *D. Msn2 binds its motifs independently in all tested contexts:* For each context, motif cooperation scores are shown as a function of the observed occupancy at 180 seconds of MNase activation. The color indicates the different libraries measured within each context. The black-circled dots are controls that include two variable positions within the same Msn2 motif. *E. Testing Msn2 Libraries in the genomic context:* A scheme illustrating CRISPR-Cas9-mediated library integration into the HO locus. *F-G: Msn2 binding in the genomic context depends on its binding sites:* Shown in (F) is the binding of Msn2 to each sequence of the *GSY2* genomically embedded library, presentation as in (C). The occupancy at the wild-type sequence in each genomically embedded library is shown in (G). *H.* Msn2 binding is independent of other TF motifs also in the genomic context: Shown are the motif pair cooperation scores as a function of the observed occupancy at 180 seconds of MNase activation in the genomic context. The black-circled dots are the above-mentioned controls that include two variable positions within the same Msn2 motif.

To test this potential influence of the broader motif context on the binding of Msn2, we selected five contexts of 2kb lengths, including a context that surrounds one of our tested regions (*GSY2*), three genomic contexts that display differential nucleosome occupancy, and a synthetic context that is expected to remain nucleosome-free (Methods; Supplementary Table 3; Figure 5B). These contexts were used to construct five new MPBA plasmids, to which eight libraries of Msn2-bound regions were integrated and tested for Msn2 binding. Notably, while these contexts affected the overall motif binding occupancies, none have led to significant motif cooperation (Figure 5C-D and Figure S6).

To further test the effect of the surrounding context, we introduced a selected set of six libraries into the genome and applied MPBA to measure Msn2 binding (Figure 5E, methods). While Msn2 still showed a strong preference for its motif, the overall binding signals to these genomically embedded libraries were somewhat lower as compared to the plasmid (Figure 5F-G). Yet, also here, none have shown motif cooperativity of significant binding effect (Figure 5H). We conclude that Msn2 retains independent motif binding across a range of different sequence contexts.

### Msn2 mutant containing only its DBD is dependent on both canonical and on similar, non-canonical motifs contributing to the binding of the full Msn2

Msn2 binding was independent of motif cooperation but was still dependent, in some cases, on motifs that better align with other TFs (Figure 6A). Therefore, Msn2 might be recruited to these regions by TFs that bind these motifs. Alternatively, Msn2 may bind those motifs directly, as suggested by their limited divergence from its canonical motif (Figure 6B). To distinguish between those possibilities, we asked whether these motifs are also bound by an Msn2 mutant containing only its DBD, which is unlikely to interact with other TFs (Figure 6C).

**Figure 6.**
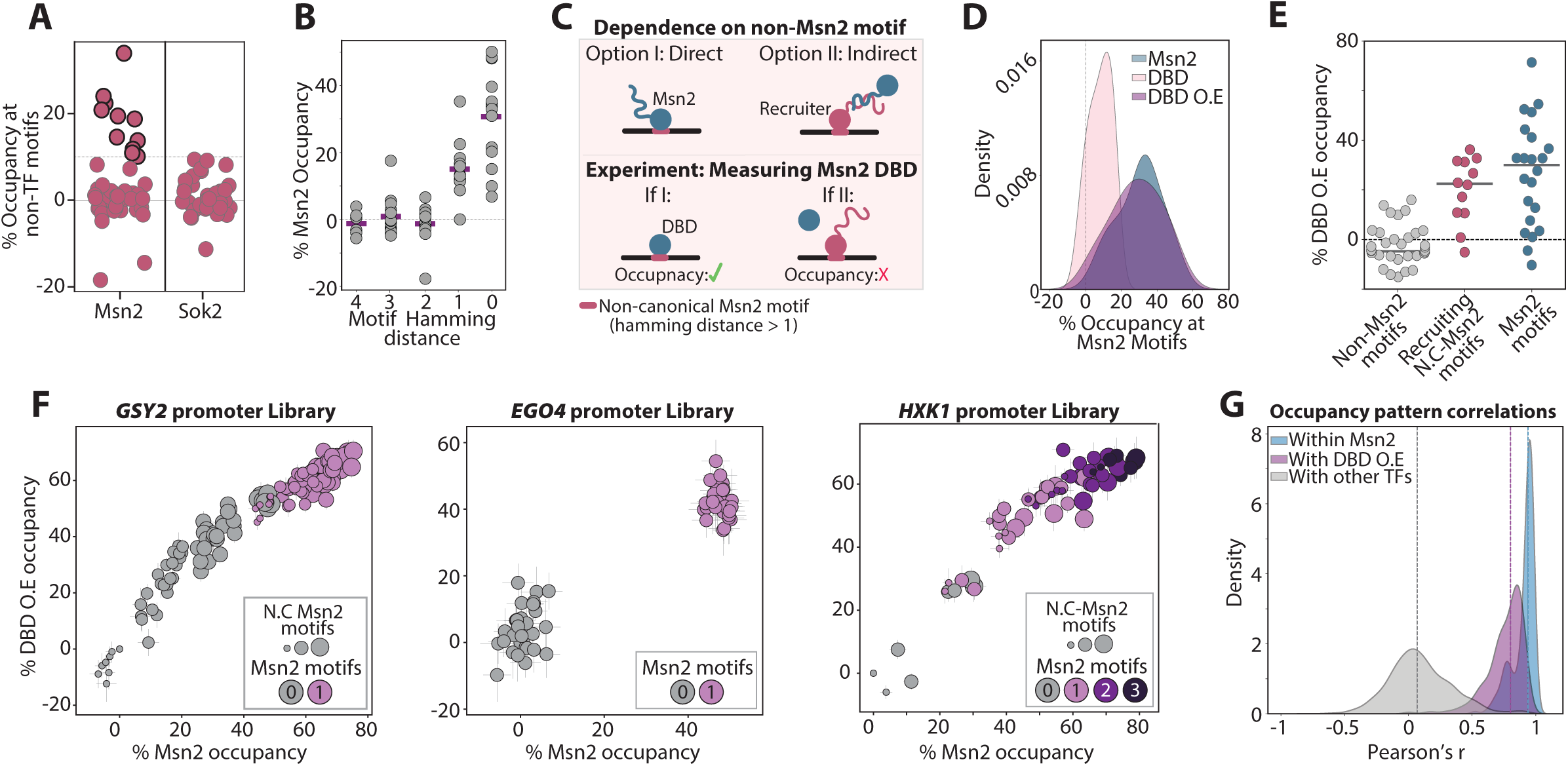
Msn2 binding to non-canonical motifs is explained by direct recognition of its DBD: *A-B. Motifs contributing to Msn2 binding are similar in sequence to its canonical motif:* The average occupancies of all Msn2 and Sok2 non-canonical motifs when all the other motifs are mutated are shown in (A; Methods). While Msn2 depends on a few non-canonical motifs, Sok2 does not depend on motifs other than its own. In (B), the Msn2 occupancies at all motifs found in the tested libraries are plotted as a function of their Hamming distance from the Msn2 motif, i.e. the number of positions in which the respective motif differs from the canonical one (AGGGG). Purple lines indicate the median Msn2 dependent occupancy for each Hamming distance. *C*. *Experimental scheme for distinction between direct and indirect binding of non-canonical Msn2 motifs:* We wished to distinguish whether Msn2 directly binds the non-canonical motifs contributing to its occupancy using its DBD (Option I, top left) or alternatively, recruited to these motifs via interaction with another TF (Option II, top right). To this end, we measured the occupancy of an Msn2 mutant containing only its DBD. We expect this mutant to lack the ability to interact with other TFs (bottom). *D. Regions outside the DBD stabilize Msn2 binding at regulatory regions:* In addition to the wild-type Msn2, we tested the binding of the DBD-only mutant across the 11 Msn2 libraries presented above. Shown are the distributions of the occupancies measured for the wild-type Msn2 and its DBD-only mutant at sequence variants in which only one Msn2 canonical motif is intact and all other motifs found within the respective region mutated, over all tested libraries. As the binding of the DBD-only variant was low, we also tested its binding upon over-expression using the strongest budding yeast promoter (*TDH3*). *E-G. Msn2 motif occupancy is explained by preferences of its DBD:* Shown in (E) are the single motif-dependent occupancies of the over-expressed Msn2 DBD-only mutant for all motifs found within the tested libraries. Sequences are classified based on the presence of a canonical Msn2 motif (AGGGG, blue), a non-canonical motif that contributes to the binding of the wild-type Msn2 (Hamming distance > 0, red; as defined in a prior experiment shown in Figure 4, based on the wild-type Msn2 at occupancy at 180 seconds of activation), and non-canonical motifs showing no contribution to the occupancy of Msn2 (gray). The direct comparisons of the measured occupancies of the wild-type and DBD-only at each individual sequence are shown in (F) for three selected libraries at 180 seconds of MNase activation (presentation as in Figure 4D). Error bar represent the SEM over repeats (5-7 repeats for each TF in each library). Shown in (G) are the distributions of correlations between occupancies at each sequence of the wild-type Msn2 and DBD-only mutant across all Msn2 libraries. The distributions of correlations between the wild-type Msn2 repeats and between Msn2 to unrelated TFs, measured across those same libraries, are also presented (see Figure 6, below). Note the tight correlation in the binding pattern between the wild-type Msn2 and its DBD-only variant.

In agreement with our previous data, the DBD-only mutant displayed weak library binding when expressed under the endogenous Msn2 promoter, limiting our ability to measure motif occupancies reliably (Figure 6D). Over-expression of the DBD overcame this apparent limited affinity, reaching high occupancies across our libraries that were comparable to the endogenously expressed Msn2 (Figure 6D). Notably, the DBD motif-binding patterns within each library correlated well with those of Msn2, and it showed similar localization not only at Msn2 motifs but also to the non-canonical, Msn2-recruiting ones (Figures 6E-G and S7). Therefore, all motifs on which Msn2 depends for binding are directly recognized by its DBD.

### Large-scale testing of motif interactions

Our analysis so far questioned a general role for motif-dependent cooperation in guiding TF localization, as the motifs contributing to the binding of the GRFs, Msn2, and Sok2 appear to have independent effects. Still, cooperative motif interactions were clearly seen in the case of Tec1 and Ste12. To examine the prevalence of cooperating motifs more broadly, we extended our analysis to dozens of additional TFs (Figures 7A-C). For this, we selected a total of 68 regions bound by multiple TFs, generated corresponding libraries including all combinations of motif mutations, and tested the binding of respective TFs to those libraries. Overall, we measured 250 TF-library pairs.

**Figure 7.**
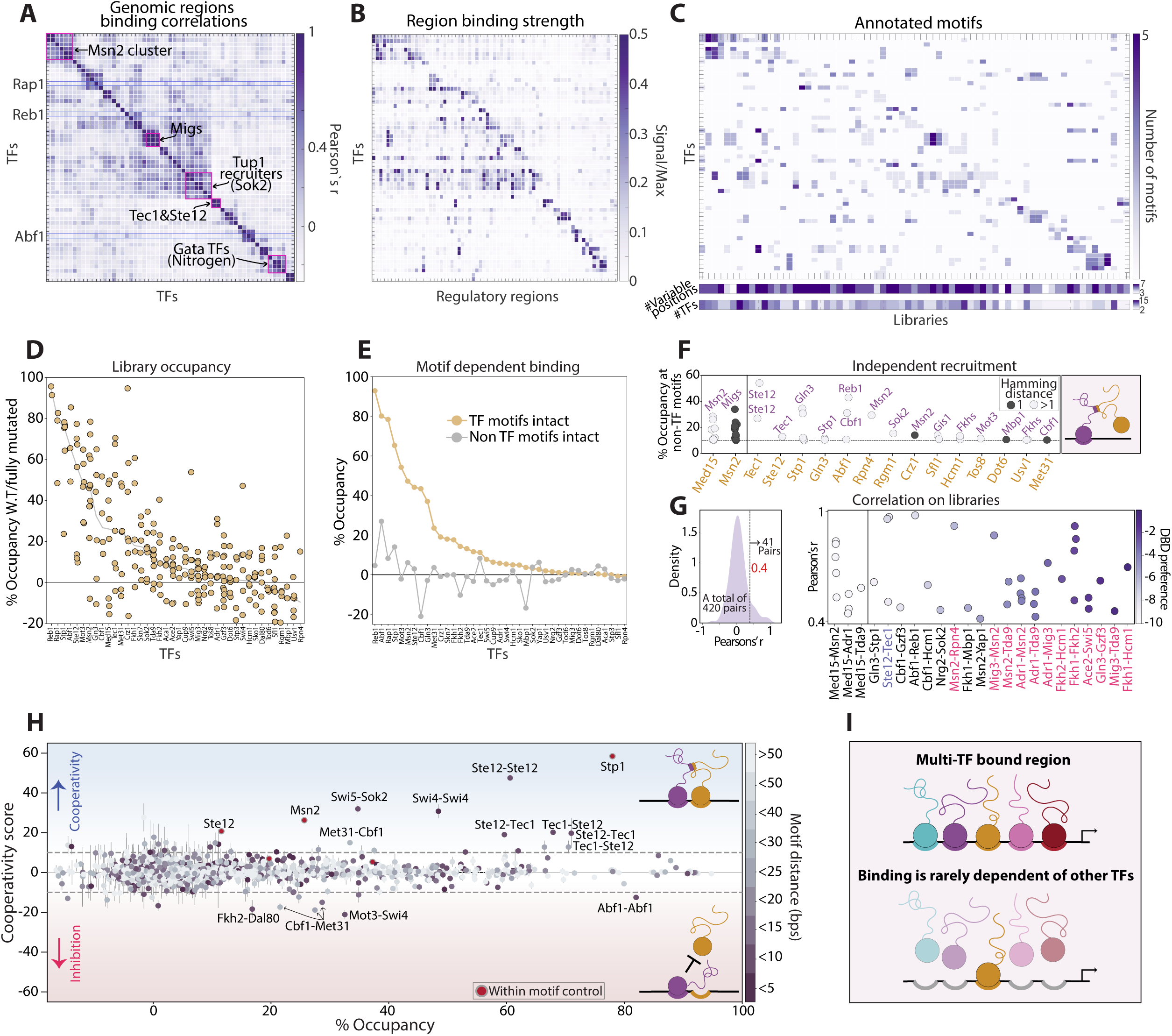
Screen for motif-dependent interactions reveals independent binding with rare cases of motif cooperation: *A-C. TFs and regulatory regions selected for screening motif interactions:* As candidates for cooperating motifs, we searched our dataset of 145 TF binding profiles for regulatory regions containing dense sets of bound motifs, as described in Figure 1A. Overall, 39 TFs, and 68 regulatory regions of the length of 164 bps were selected. Shown in (A) is the similarity (correlation) of the genomic binding profiles of the selected TFs, measured across the selected regions. Note the high binding similarity of the interacting factors Ste12-Tec1, as well as the clusters of TFs that bind at promoters overlapping with Msn2 or Sok2. Shown in (B) are the TF binding signals themselves, presented as the ratio between the total signal received for each TF on each region to the maximal signal received by the same TF on a similar-sized region when examining the entire genome (Methods). The number of respective TF motifs in each selected region is shown in (C). The overall number of motifs within each regulatory region, as well as the overall number of TFs predicted to bind these same motifs are indicated (C, bottom stripes) *D-E. Motif-dependent TF occupancy across regulatory regions:* TF occupancies at each regulatory sequence are shown in (D), calculated using the abundance of intact vs. fully mutated sequences. TFs are ordered by the median occupancies of their respective regulatory regions (gray line). The self-preferred motif dependent region occupancy, measured by comparing the average occupancy at sequences containing intact vs. mutated canonical motifs of each TF, are shown in (E, orange line; Methods). Average region occupancies depending on non-canonical motifs are also plotted (gray line). *F-G One sided recruitment is limited:* TF occupancy could depend not only on self canonical motifs, but also on motifs associated with other TFs, potentially indicating recruitment as illustrated in the scheme shown in (F). To estimate this dependency, we measured the contribution to occupancy of each non-canonical motif when the canonical motifs are mutated (Methods). Shown on the left are cases where such dependency was observed (>10% contribution to occupancy). The tested TFs are indicated at the bottom (orange), and the TF-associated with the recruiting motif is indicated within the plot (purple). The co-activator Med15 is known to be recruited by Msn2, and was tested on 7 of the Msn2-selected regulatory regions as a control. Msn2 dependencies are the same as shown in Figure 5A for non-canonical motifs contributing more than 10% to its occupancy. The Hamming distance between the affecting motif and the TF-canonical motif is color-coded. As an additional measure for dependency, we calculated the correlation in the occupancy levels at each sequence between different TFs tested on the same regions. Most TFs displayed distinct patterns, consistent with differential motif-dependencies (G, left). The small number of cases showing high correlations (Pearson’s r > 0.4) are highlighted (G, right), and include the Med15 high correlation with Msn2 in 5 out of 7 testes libraries. Each dot corresponds to one library, tested for binding by both TFs indicated on the x-axis. Color code indicates the Euclidean distances between the respective TF DBDs preferred motifs (Methods). TF pairs from the same DBD family are indicated in pink. Note the high correlation calculated for Ste12 and Tec1 and the high similarity in DBD preferences of most detected TF pairs. *H. Cooperation among motif pairs is rare:* Shown are the cooperation scores amongst all 1917 motif pairs containing at least one canonical motif of the TFs measured in our screen, as a function of the observed occupancy over all variants containing the motif pairs intact. Colors indicate the distance between the two motifs within each pair two motifs. Highlighted are pairs for which the interaction (cooperation or inhibition) was high, where the TF measured is indicated on the left and the TF associated with the second motif found on the right. Control cases carrying two variable positions found within the same motif are marked in red. Note the small fraction of cooperating motif pairs; only 36 of the 1917 (<2%) showed cooperativity scores higher than 10%. Out of 606 motif pairs contributing > 10% to TF occupancy, only 17 show cooperation (2.8%). *I. Summary scheme:* Based on our screen we conclude that TF binding is well explained by the additive contributions of individual motifs. Thus, motif-dependent cooperativity is rare, refuting its general role in increasing TF genomic binding specificity.

The estimated occupancies at intact regulatory regions varied between TFs. High occupancies that consistently reached ~60-90% were seen for Stp1, Mot3, and Cbf1, as well as for the GRFs discussed above (Figure 7D). Further, in 33/39 of the TFs tested, the average occupancy at their own motif exceeded that of other TF motifs (Figure 7E). In some cases, additional, non-canonical motifs contributed to binding, and those included sequence-similar motifs (e.g., The Met31 and the Cbf1) but also non-similar ones, which may suggest recruitment (e.g., Rgm1 by Sok2 or Gln3 by Stp1; Figure 7F). To verify this, we compared the correlation in motif effects of different TFs measured across the same library. For this, we used Med15, an Msn2-recruited coactivator (44), as a control, measuring its binding across the Msn2 libraries (Figure 7F-G). Notably, only ~10% of the TF pairs showed similar motif effects (Pearson’s r>0.4), and these included the Msn2 and Med15 control, as well as paralogs whose DBDs indeed bind the same motif. Still, eight TF pairs localized to similar motifs despite having distinct DBDs, and those included the known Ste12-Tec1 and Gln3-Stp1, noted above.

Next, we defined the cooperation score for all 1917 pairs, including at least one motif of the tested TF, available in our dataset (Figures 7H and S8A). Apart from Ste12-Tec1 discussed above, a few more cooperating motif pairs were detected. These include the Swi5-Sok2 motifs, whose cooperation increased Swi5 binding by 32% at the *RSM7* promoter, and two Swi4 motifs, whose cooperation increased Swi4 binding by 31% at the *CAP1* promoter. Notably, all of these pairs were located in high proximity to one another (Figures 7H and S8B). Those cooperating motif pairs, however, were the exception. Out of 606 pairs showing > 10% contribution to TF occupancy, 589 (97%) were independent of motif cooperation. Therefore, at the global scale, TF localization to their genomic binding sites is largely invariant to the presence of nearby motifs (Figure 7I).

## Discussion

Studies in cell-free systems defined the mechanisms through which DNA binding domains (DBDs) recognize specific DNA motifs. This binding of TF DBDs to their preferred motifs also occurs inside cells, but for this, the TFs need to first detect their binding sites amidst the vast number of similar sequences. In this study, we systematically examined the widely assumed role of motif combinatorics in distinguishing TF binding sites from other motif occurrences that remain unbound. Our main finding is that motif cooperativity is rare. Rather, we find that in most cases, when multiple motifs contribute to the binding of a TF, they act additively so that the overall occupancy is well explained by the independent contribution of each motif.

We chose Msn2 and Sok2 as the focus of our analysis, as both bind promoters that contain multiple additional binding sites of other TFs, and both are directed to those regions through long intrinsically disordered regions (IDRs) located outside their DBDs. Still, of the 164 motif pairs within the 17 tested regulatory regions, only one exhibited cooperative binding, and even then, the contribution to the already strong binding was marginal. Instead, multiple motifs appeared to contribute to binding in an additive manner.

Notably, this lack of cooperative interactions occurred despite the significant influence of the bound TFs on nucleosomes. The Binding of Msn2, for example, has led to the loss of nucleosomes at all tested regions. Independent motif binding also remained when integrating the tested regulatory region within different sequence contexts associated with distinct nucleosome architecture. Further, extending our plasmid-based assay to enable genomic integration confirmed the lack of cooperativity also under this condition. Together, these results largely refute the role of motif combinatorics in explaining the binding specificity of Sok2 and Msn2.

Through further analysis, in which we tested the binding of an Msn2 mutant containing only its DBD, we also refuted the possibility that Msn2 is recruited to its target promoters through interactions with other TFs. While this possibility does not explain its binding specificity, it could account for the localization of the mutant lacking the DBD to those regions(19). It is notable that while expressed under the native Msn2 promoter, the DBD-only mutant showed poor overall binding to the tested libraries, consistent with the known role of regions located outside of the DBD in stabilizing the binding to those promoters.

Our finding that TFs bind to individual motifs, irrespectively of proximal sites, challenges prevailing thinking attributing binding specificity to combinatorial motif usage. This leaves open the question of what distinguishes bound motifs from those remaining unoccupied. Limited chromatin accessibility may prevent TFs from binding certain motifs, yet this provides a partial solution only and does not explain the prominent role of non-DBD regions in directing TF binding. Whether specificity arises through interactions with chromatin or the DNA itself remains to be investigated.

While being primarily interested in motif-dependent cooperation, we also obtained estimates of TF occupancies at the various regulatory regions, as well as at their individual motifs within those regions. Those occupancies varied greatly between TFs and between the selected regulatory regions, ranging from cases in which the region appeared to be bound in >90% of cells to cases where binding was too low to be detected. Msn2 was amongst the strongest bound TFs, as well as other TFs that have not yet received much experimental attention, such as Stp1 and Mot3. A large fraction of TFs showed intermediate occupancies of <20-30%, even at their top targets. Although the binding profiles used to select the multi-TF bound regulatory regions were measured under the same conditions used in this study (OD_600_=4), these low occupancies could reflect their low activation or abundance in these conditions. Occupancies increased with MNase activation times, indicating an increased fraction of cells in which the respective TF was bound. While we did not explore this systematically, the rates at which those occupancies increased differed between TFs and regions, which may be used to quantify properties of TF-target search kinetics, including the respective association and dissociation rates.

Our study was made possible due to a novel method we developed that enables measuring TF binding to thousands of designed DNA sequences in parallel. Such measurements were crucial in moving beyond merely correlating motif proximities or indirectly measuring the downstream effects of TF interactions to defining the contribution of combinatorics to TF binding. While considerable effort has been made to achieve these direct measurements (45), MPBA accomplishes this and enables a temporally controlled readout of instantaneous binding events. Being sequencing-based, our method is rapid, quantitative, and relatively simple to implement. It is also expression-independent and has the ability to monitor instantaneous binding events upon MNase activation. By this, our method offers a promising new platform for future applications requiring direct *in vivo* analysis of DNA-protein interactions.

## Data availability

Further information and requests for resources and reagents should be directed to and will be fulfilled by the Lead Contact, Dr. Naama Barkai. All strains used in this study are available by direct request to the lead contact without any further restrictions

Code and processed data are available in [GitHub](https://github.com/sagieb/MPBA). All raw sequencing data generated in this study have been deposited in GEO under accession number: GSE234455.

## References

1. Wunderlich, Z. and Mirny, L.A. (2009) Different gene regulation strategies revealed by analysis of binding motifs. Trends Genet, 25, 434–440.

2. Wang, J., Zhuang, J., Iyer, S., Lin, X.Y., Whitfield, T.W., Greven, M.C., Pierce, B.G., Dong, X., Kundaje, A., Cheng, Y., et al. (2012) Sequence features and chromatin structure around the genomic regions bound by 119 human transcription factors. Genome Res, 22, 1798–1812.

3. Jana, T., Brodsky, S. and Barkai, N. (2021) Speed-Specificity Trade-Offs in the Transcription Factors Search for Their Genomic Binding Sites. Trends Genet, 37, 421–432.

4. Neph, S. (2012) An expansive human regulatory lexicon encoded in transcription factor footprints. Nature, 489, 83–90.

5. Inukai, S., Kock, K.H. and Bulyk, M.L. (2017) Transcription factor-DNA binding: beyond binding site motifs. Curr Opin Genet Dev, 43, 110–119.

6. Dror, I., Rohs, R. and Mandel-Gutfreund, Y. (2016) How motif environment influences transcription factor search dynamics: Finding a needle in a haystack. Bioessays, 38, 605– 612.

7. Reiter, F., Wienerroither, S. and Stark, A. (2017) Combinatorial function of transcription factors and cofactors. Curr Opin Genet Dev, 43, 73–81.

8. Morgunova, E. and Taipale, J. (2017) Structural perspective of cooperative transcription factor binding. Curr Opin Struct Biol, 47, 1–8.

9. Vierstra, J. (2020) Global reference mapping of human transcription factor footprints. Nature, 583, 729–736.

10. Arnosti, D.N. and Kulkarni, M.M. (2005) Transcriptional enhancers: Intelligent enhanceosomes or flexible billboards? J Cell Biochem, 94, 890–898.

11. Lin, Y.-S. (1990) How different eukaryotic transcriptional activators can cooperate promiscuously. Nature, 345, 359–361.

12. Vernot, B. (2012) Personal and population genomics of human regulatory variation. Genome Res, 22, 1689–1697.

13. Rastegar, S., Hess, I., Dickmeis, T., Nicod, J.C., Ertzer, R., Hadzhiev, Y., Thies, W.G., Scherer, G. and Strähle, U. (2008) The words of the regulatory code are arranged in a variable manner in highly conserved enhancers. Dev Biol, 318, 366–377.

14. Levo, M., Avnit-Sagi, T., Lotan-Pompan, M., Kalma, Y., Weinberger, A., Yakhini, Z. and Segal, E. (2017) Systematic Investigation of Transcription Factor Activity in the Context of Chromatin Using Massively Parallel Binding and Expression Assays. Mol Cell, 65, 604–617.e6.

15. Deplancke, B., Alpern, D. and Gardeux, V. (2016) The Genetics of Transcription Factor DNA Binding Variation. Cell, 166, 538–554.

16. Kim, S. and Wysocka, J. (2023) Deciphering the multi-scale, quantitative cis-regulatory code. Mol Cell, 83, 373–392.

17. Mirny, L.A. (2010) Nucleosome-mediated cooperativity between transcription factors. Proc Natl Acad Sci U S A, 107, 22534–22539.

18. Adams, C.C. and Workman, J.L. (1995) Binding of disparate transcriptional activators to nucleosomal DNA is inherently cooperative. Mol Cell Biol, 15, 1405–1421.

19. Brodsky, S., Jana, T., Mittelman, K., Chapal, M., Kumar, D.K., Carmi, M. and Barkai, N. (2020) Intrinsically Disordered Regions Direct Transcription Factor In Vivo Binding Specificity. Mol Cell, 79, 459–471.e4.

20. Gera, T. (2022) Evolution of binding preferences among whole-genome duplicated transcription factors. Elife.

21. Kumar, D.K. (2023) Complementary strategies for directing in vivo transcription factor binding through DNA binding domains and intrinsically disordered regions. Mol Cell.

22. Gietz, R.D., Schiestl, R.H., Willems, A.R. and Woods, R.A. (1995) Studies on the transformation of intact yeast cells by the LiAc/SS-DNA/PEG procedure. Yeast, 11, 355–360.

23. Picelli, S., Björklund, Å.K., Reinius, B., Sagasser, S., Winberg, G. and Sandberg, R. (2014) Tn5 transposase and tagmentation procedures for massively scaled sequencing projects. Genome Res, 24, 2033.

24. Köster, J., Mölder, F., Jablonski, K.P., Letcher, B., Hall, M.B., Tomkins-Tinch, C.H., Sochat, V., Forster, J., Lee, S., Twardziok, S.O., et al. (2021) Sustainable data analysis with Snakemake. F1000Res, 10.

25. Marcel Martin (2011) Cutadapt removes adapter sequences from high-throughput sequencing reads. EMBnet J, 17.

26. Lindgreen, S. (2012) AdapterRemoval: easy cleaning of next-generation sequencing reads. BMC Res Notes, 5, 337.

27. Lupo, O., Kumar, D.K., Livne, R., Chappleboim, M., Levy, I. and Barkai, N. (2023) The architecture of binding cooperativity between densely bound transcription factors. Cell Syst, 14, 732–745.e5.

28. Weirauch, M.T., Yang, A., Albu, M., Cote, A.G., Montenegro-Montero, A., Drewe, P., Najafabadi, H.S., Lambert, S.A., Mann, I., Cook, K., et al. (2014) Determination and inference of eukaryotic transcription factor sequence specificity. Cell, 158, 1431–1443.

29. Langmead, B. and Salzberg, S.L. (2012) Fast gapped-read alignment with Bowtie 2. Nat Methods, 9, 357–359.

30. Langmead, B., Wilks, C., Antonescu, V. and Charles, R. (2019) Scaling read aligners to hundreds of threads on general-purpose processors. Bioinformatics, 35, 421–432.

31. Quinlan, A.R. and Hall, I.M. (2010) BEDTools: a flexible suite of utilities for comparing genomic features. Bioinformatics, 26, 841–842.

32. Kaplan, N., Moore, I.K., Fondufe-Mittendorf, Y., Gossett, A.J., Tillo, D., Field, Y., LeProust, E.M., Hughes, T.R., Lieb, J.D., Widom, J., et al. (2009) The DNA-encoded nucleosome organization of a eukaryotic genome. Nature, 458, 362–366.

33. Zhao, H., Sun, Z., Wang, J., Huang, H., Kocher, J.P. and Wang, L. (2014) CrossMap: a versatile tool for coordinate conversion between genome assemblies. Bioinformatics, 30, 1006– 1007.

34. Sharon, E. (2012) Inferring gene regulatory logic from high-throughput measurements of thousands of systematically designed promoters. Nat Biotechnol, 30, 521–530.

35. Haberle, V. and Lenhard, B. (2012) Dissecting genomic regulatory elements in vivo. Nat Biotechnol, 30, 504–506.

36. Patwardhan, R.P. (2012) Massively parallel functional dissection of mammalian enhancers in vivo. Nat Biotechnol, 30, 265–270.

37. Melnikov, A. (2012) Systematic dissection and optimization of inducible enhancers in human cells using a massively parallel reporter assay. Nat Biotechnol, 30, 271–277.

38. Boer, C.G. (2020) Deciphering eukaryotic gene-regulatory logic with 100 million random promoters. Nat Biotechnol, 38, 56–65.

39. Schmid, M., Durussel, T. and Laemmli, U.K. (2004) ChIC and ChEC; genomic mapping of chromatin proteins. Mol Cell, 16, 147–157.

40. Fourel, G. (2002) General Regulatory Factors (GRFs) as Genome Partitioners*. Journal of Biological Chemistry, 277, 41736–41743.

41. Chasman, D.I. (1990) A yeast protein that influences the chromatin structure of UASG and functions as a powerful auxiliary gene activator. Genes Dev, 4, 503–514.

42. Madhani, H.D. and Fink, G.R. (1997) Combinatorial control required for the specificity of yeast MAPK signaling. Science (1979), 275, 1314–1317.

43. Hoi, J.W.S. and Dumas, B. (2010) Ste12 and Ste12-Like Proteins, Fungal Transcription Factors Regulating Development and Pathogenicity. Eukaryot Cell, 9, 480–485.

44. Sadeh, A., Baran, D., Volokh, M. and Aharoni, A. (2012) Conserved motifs in the Msn2-activating domain are important for Msn2-mediated yeast stress response. J Cell Sci, 125, 3333–3342.

45. Liu, J., Shively, C.A. and Mitra, R.D. (2020) Quantitative analysis of transcription factor binding and expression using calling cards reporter arrays. Nucleic Acids Res, 48, e50–e50.

