## Supplemental Figures for "Massively Parallel Binding Assay (MPBA) reveals limited transcription factor binding cooperativity, challenging models of specificity"

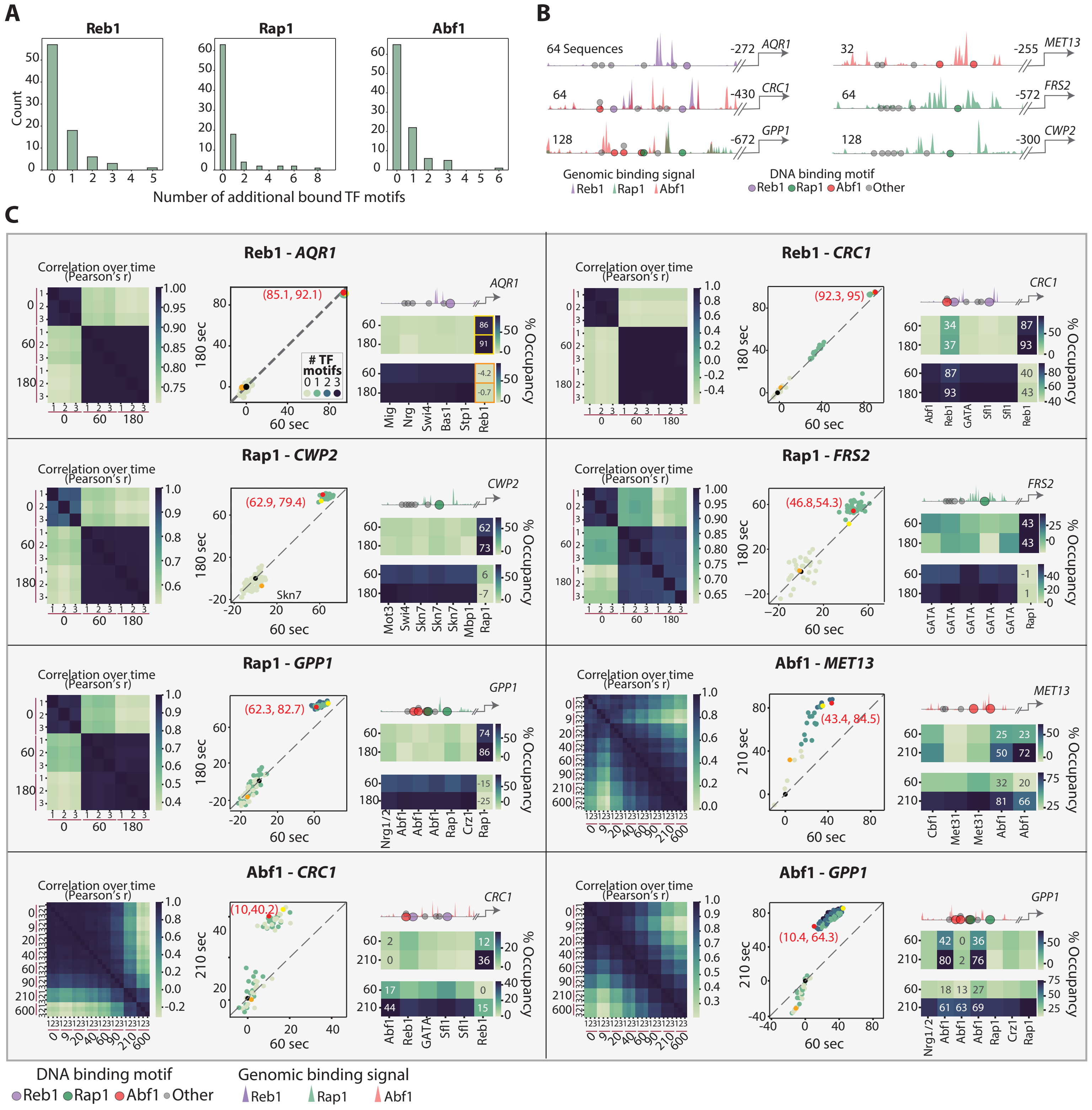
A
Reb1
Rap1
Abf1
Count
Number of additional bound TF motifs
B
64 Sequences
-272 AQR1
64
-430 CRC1
128
-672 GPP1
32
-255 MET13
64
-572 FRS2
128
-300 CWP2
Genomic binding signal
Reb1 Rap1 Abf1
DNA binding motif
Reb1 Rap1 Abf1 Other
C
Reb1 - AQR1
Rap1 - CWP2
Rap1 - GPP1
Abf1 - CRC1
Reb1 - CRC1
Rap1 - FRS2
Abf1 - MET13
Abf1 - GPP1
Correlation over time (Pearson's r)
180 sec
210 sec
60 sec
% Occupancy
### TF motifs
(85.1, 92.1)
(92.3, 95)
(62.9, 79.4)
(46.8,54.3)
(62.3, 82.7)
(43.4, 84.5)
(10,40.2)
(10.4, 64.3)
Skn7
Mig Nrg Swi4 Bas1 Stp1 Reb1
Abf1 Reb1 GATA Sfl1 Sfl1 Reb1
Mot3 Swi4 Skn7 Skn7 Skn7 Mbp1 Rap1
GATA GATA GATA GATA GATA Rap1
Nrg1/2 Abf1 Abf1 Abf1 Rap1 Rap1
Cbf1 Met31 Met31 Abf1 Abf1
Abf1 Reb1 GATA Sfl1 Sfl1 Reb1
Nrg1/2 Abf1 Abf1 Abf1 Rap1 Crz1 Rap1
DNA binding motif
Reb1 Rap1 Abf1 Other
Genomic binding signal
Reb1 Rap1 Abf1

**Figure S1. MPBA shows high reproducibility, revealing GRFs dependency on their own motifs.**

(A) *The GRFs bind promoters depleted of other TFs:* The top bound promoters of each GRF were searched for bound GRF motifs found in proximity to another TF motif enriched in binding signal of the respective TF (distance <90 bps, Methods). Shown are the number of additional bound TF motifs found in proximity to all identified GRF bound motifs. Note that in most cases, there are no additional TFs bound at sites found in proximity to the GRF binding site.

(B) *Regulatory regions selected for measuring motif-dependent GRF binding:* We selected six regulatory regions bound by at least one of the three GRFs. Shown are the ChEC-seq binding signals across the selected regions, GRFs are indicated by colors (Methods), where the signal of each GRF is normalized to its maximal signal received in this region. Distances of the selected regions from the TSS of the nearest genes are noted. The locations of all identified TF motifs are displayed as circles, while motifs corresponding to a GRF are shown in colors. Based on these motifs, six libraries of combinatorial mutations were generated. The number of sequences in each library (32-128) is also indicated. These libraries were tested for the binding of the relevant GRFs.

(C) *GRFs reach high occupancy at individual motifs and exhibit high reproducibility:* In each panel, shown on the left is the sequence abundance correlation for the indicated library bound by the indicated TF in each time point and for all technical repeats (3 per time point). The middle of each panel shows the occupancy of the GRF at each library sequence (dots) at the indicated MNase activation time-points. The number of intact GRF motifs in each library sequence is color-coded. The dots representing the fully intact sequence (and its corresponding values) and the fully mutated variant are indicated in red and black respectively. Orange color represents the variant in which the motifs of the tested GRF are intact and those of other TFs are mutated, and yellow color marks the variant in which the tested GRF motifs are mutated while other TF motifs are intact. On the right of each panel is the relevant promoter (top; presentation as in S1A) and two heatmaps: The upper one shows the GRF occupancy at sequences in which all positions are mutated except for a single one, represented by each column, and the lower heatmap shows the GRF occupancy at sequences in which all positions are intact except for a single one.

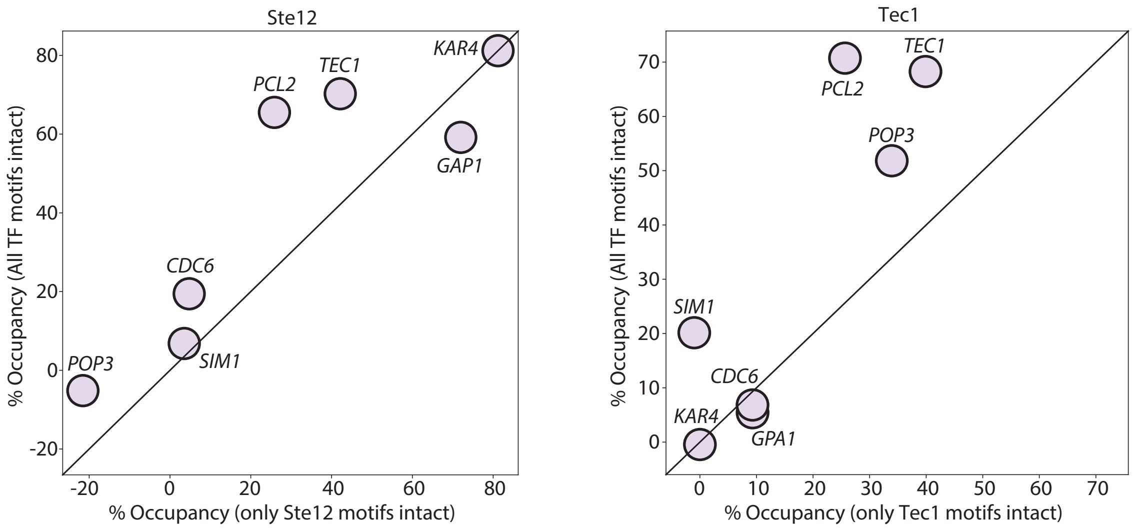

Ste12
% Occupancy (All TF motifs intact)
% Occupancy (only Ste12 motifs intact)
-20
0
20
40
60
80
KAR4
TEC1
PCL2
GAP1
CDC6
SIM1
POP3
Tec1
% Occupancy (All TF motifs intact)
% Occupancy (only Tec1 motifs intact)
0
10
20
30
40
50
60
70
PCL2
TEC1
POP3
SIM1
CDC6
KAR4
GPA1

**Figure S2. Tec1 and Ste12 reach high occupancy and depend on the motifs of one another.**

Shown is the occupancy of Tec1 and Ste12 at each intact regulatory region, relative to the fully mutated sequence, as a function of the occupancy of the same region when only the motifs of the tested TF are intact. Note the high dependency of both Tec1 and Ste12 on motifs other than their own in both *PCL2* and *TEC1* promoter libraries, suggesting their cooperation.

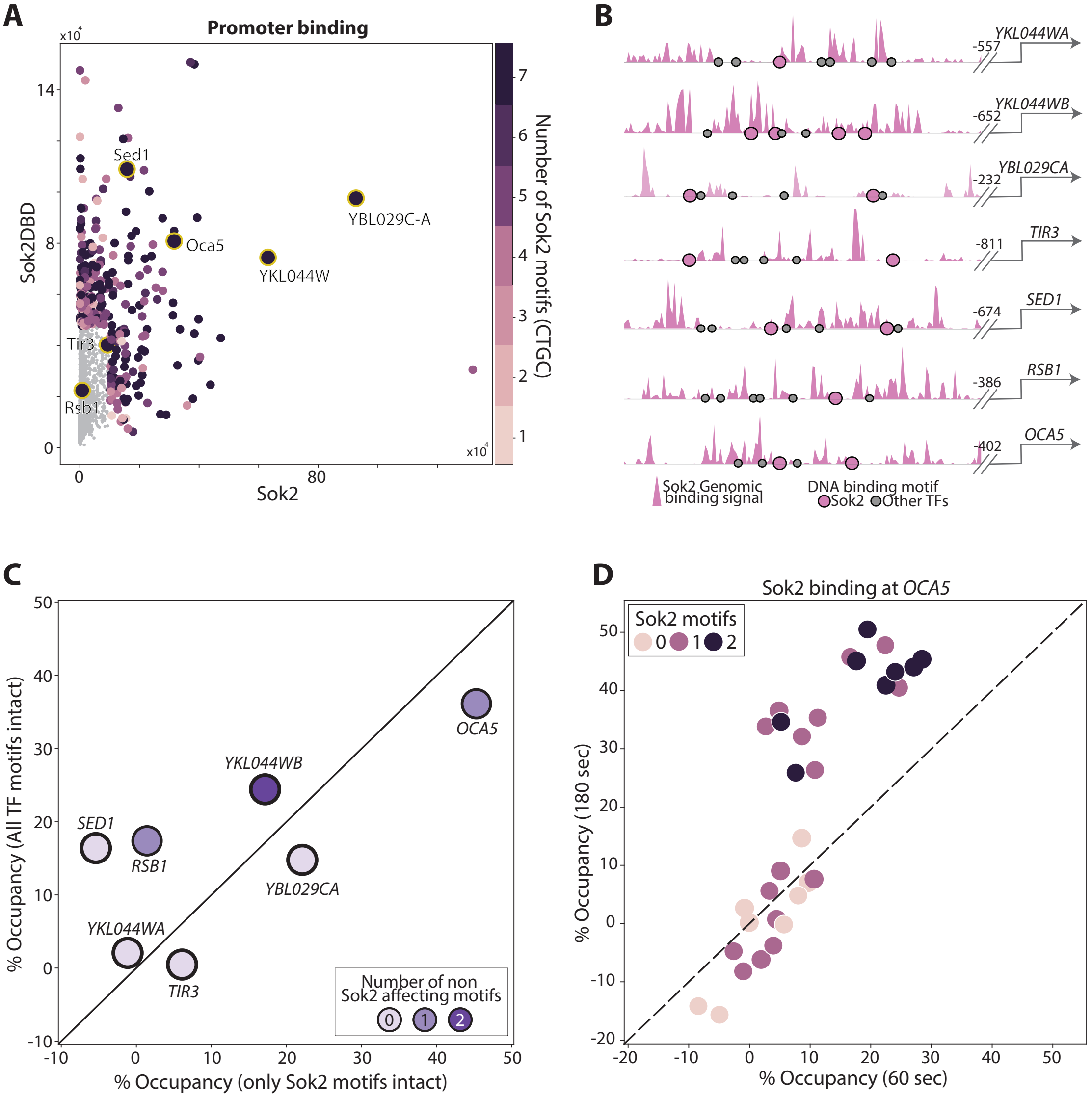
A
Promoter binding
x10^4
14
8
0
Sok2DBD
Sed1
YBL029C-A
Oca5
YKL044W
Tir3
Rsb1
0
80
Sok2
x10^4
Number of Sok2 motifs (CTGC)
7
6
5
4
3
2
1
B
YKL044WA
-557
YKL044WB
-652
YBL029CA
-232
TIR3
-811
SED1
-674
RSB1
-386
OCA5
-402
Sok2 Genomic binding signal
DNA binding motif
Sok2
Other TFs
C
% Occupancy (All TF motifs intact)
% Occupancy (only Sok2 motifs intact)
OCA5
YKL044WB
SED1
RSB1
YBL029CA
YKL044WA
TIR3
Number of non Sok2 affecting motifs
0
1
2
50
40
30
20
10
0
-10
D
Sok2 binding at OCA5
Sok2 motifs
0
1
2
% Occupancy (180 sec)
% Occupancy (60 sec)
-20
-10
0
10
20
30
40
50

**Figure S3. Sok2 binds mostly independently of other TFs.**

(A-B) *Selection of regulatory regions:* Shown in (A) are the sum of signals received on each promoter in the genome, comparing Sok2 to a mutant containing only its DNA Binding domain (DBD). The number of Sok2 motifs found in each promoter sequence is indicated by color. The chosen Sok2 bound regulatory regions are circled in yellow and shown in (B; presentation as in S1B).

(C-D) *Sok2 depends mainly on its own motifs for binding:* Shown in (C) is the occupancy of Sok2 at each intact regulatory region, relative to the fully mutated sequence, as a function of the occupancy of the same region when only the canonical Sok2 motifs are intact. The occupancy of each sequence of the library based on the *OCA5* regulatory region is shown in (D). Color indicates the number of intact Sok2 motifs in each of the library sequences.

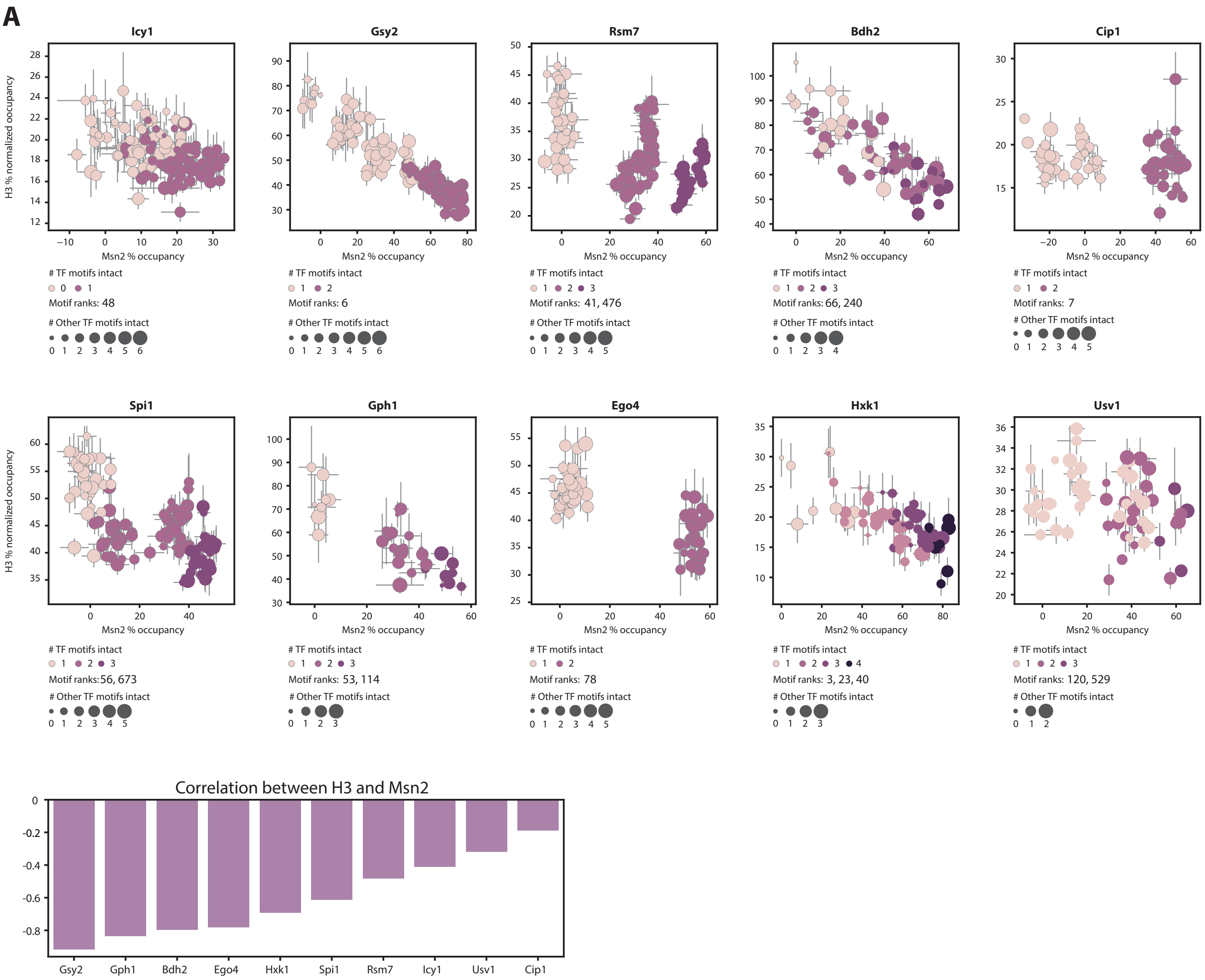
A
Icy1
Gsy2
Rsm7
Bdh2
Cip1
H3 % normalized occupancy
Msn2 % occupancy
### TF motifs intact
0 1
Motif ranks: 48
### Other TF motifs intact
0 1 2 3 4 5 6
1 2
Motif ranks: 6
1 2 3
Motif ranks: 41, 476
0 1 2 3 4 5
Motif ranks: 66, 240
0 1 2 3 4
1 2
Motif ranks: 7
Spi1
Gph1
Ego4
Hxk1
Usv1
Motif ranks:56, 673
Motif ranks: 53, 114
Motif ranks: 78
1 2 3 4
Motif ranks: 3, 23, 40
Motif ranks: 120, 529
B
Correlation between H3 and Msn2
0
-0.2
-0.4
-0.6
-0.8
Gsy2
Gph1
Bdh2
Ego4
Hxk1
Spi1
Rsm7
Icy1
Usv1
Cip1

**Figure S4. Msn2 binding causes a proportional eviction of nucleosomes in all tested libraries.**

(A-B) Shown in (A), for each of the indicated libraries, is the H3 normalized occupancy (Methods), as a function of the occupancy of Msn2. Each dot represents a library variant, colored by the number of intact Msn2 motifs it contains. The size indicates the number of other TF intact motifs found in each sequence. Also indicated are the ranks of the Msn2 motifs found in each library, according to the collected genomic ChEC-seq signal. Displayed in (B) are the correlations of the scatterplots shown in (A).

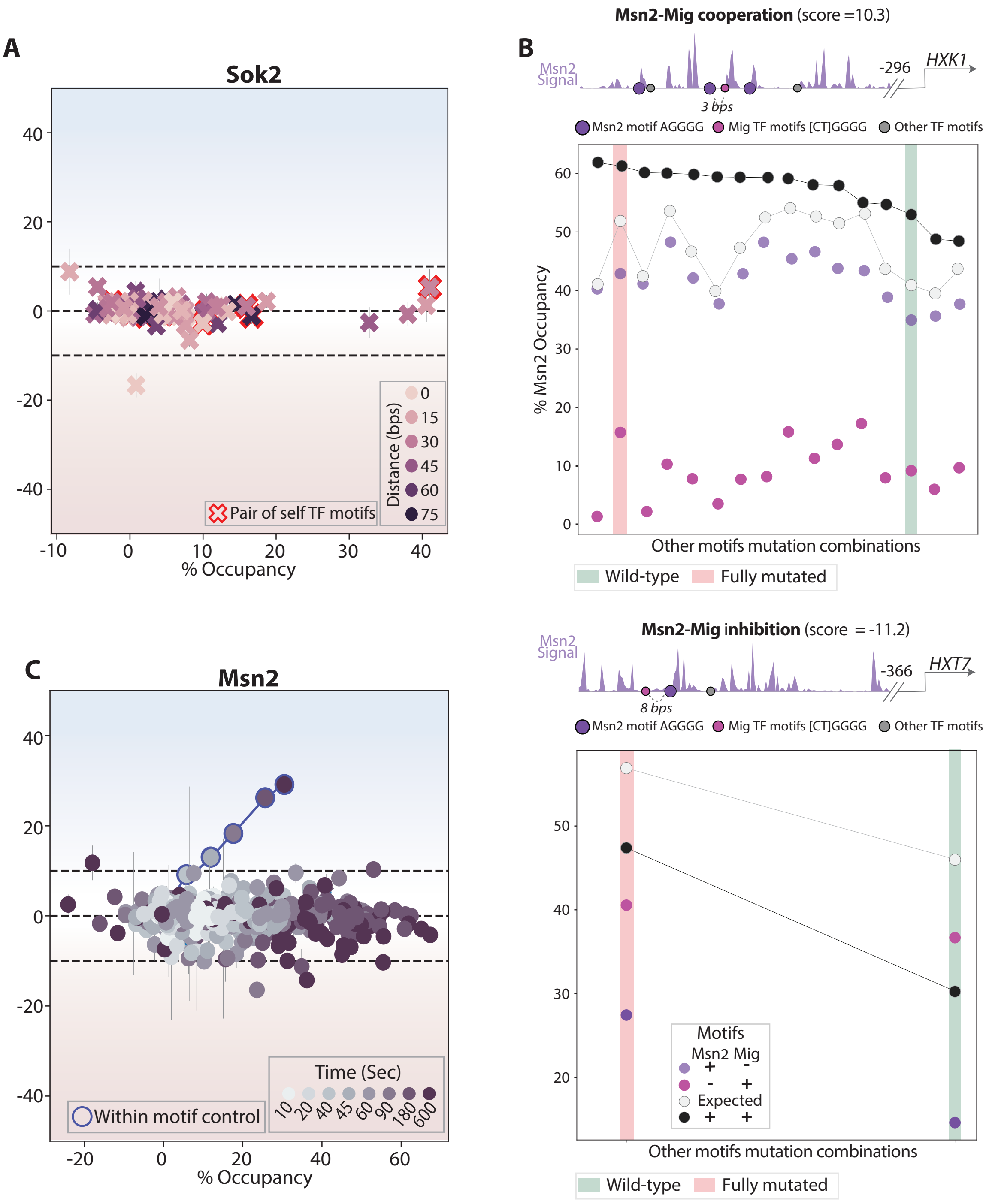
A
Sok2
% Occupancy
Distance (bps)
0
15
30
45
60
75
Pair of self TF motifs
B
Msn2-Mig cooperation (score =10.3)
Msn2 Signal
-296
HXK1
3 bps
Msn2 motif AGGGG
Mig TF motifs [CT]GGGG
Other TF motifs
% Msn2 Occupancy
Other motifs mutation combinations
Wild-type
Fully mutated
Msn2-Mig inhibition (score = -11.2)
Msn2 Signal
-366
HXT7
8 bps
Msn2 motif AGGGG
Mig TF motifs [CT]GGGG
Other TF motifs
Motifs
Msn2 Mig
+ -
- +
Expected
+ +
Other motifs mutation combinations
Wild-type
Fully mutated
C
Msn2
% Occupancy
Time (Sec)
10 20 40 45 60 90 180 600
Within motif control

**Figure S5. Msn2 and Sok2 show no motif cooperativity.**

(A) *Msn2 additive binding is independent of MNase activation time:* Msn2 motif cooperation was tested along an MNase activation time course. Shown are the scores as a function of the observed occupancy of all motif pairs containing at least one Msn2 canonical motif within the Msn2-testing libraries, with time-points indicated by color. Note that the occupancy increases with activation time, but, with the exception of the control, cooperation scores remain low.

(B) *Rare cases of cooperativity and interruption of Msn2:* Shown are the positive (left) and inhibitory (right) effects between two pairs of Msn2-Mig TFs motifs on the Msn2 occupancy in the indicated libraries (top, presentation as in Figure S1B) for all combinations of non-pair motifs. For example, the *HXK1* library (left) contains a total of 6 motifs, therefore, we can test the effect of the pair of interest in 16 different combinations of the remaining 4 motifs (x axis). Within each such context, we compared the occupancy of three mutation combinations in the motif pair ($Msn2_{+}$-$Mig3_{-}$; $Msn2_{-}$-$Mig3_{+}$; $Msn2_{+}$-$Mig3_{+}$). Shown are the occupancies of each of those three sequences within each context of other motif combination (y axis), sorted by the observed occupancy when both motifs are intact. The respective expected occupancy of both motifs, assuming independence, is also shown (white). Fully intact and fully mutated contexts are marked by color (green and pink respectively).

(C) *Sok2 binding is independent of motif cooperation:* Motif cooperation scores are shown as a function of the observed occupancy for all 67 motif pairs containing at least one Sok2 motif within the Sok2-testing libraries. Color-code indicates the distance between the tested motifs of each pair.

A

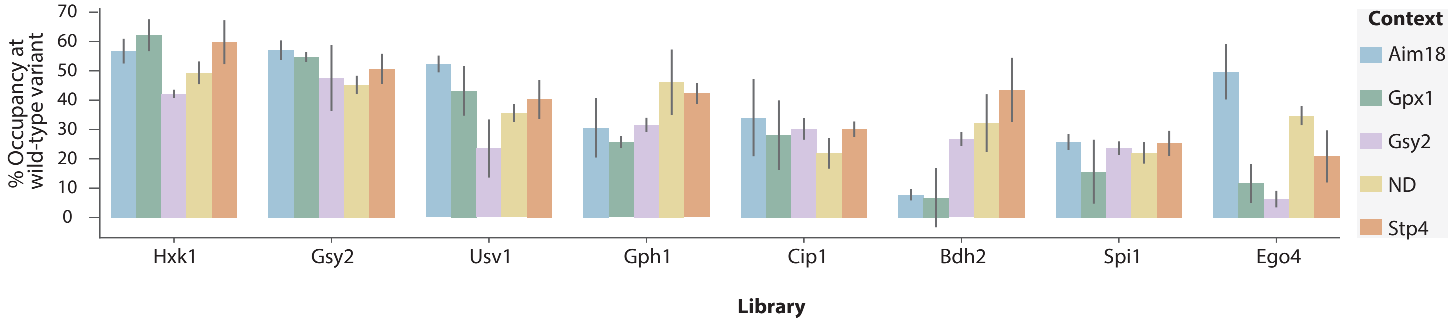
% Occupancy at wild-type variant
70
60
50
40
30
20
10
0
Hxk1
Gsy2
Usv1
Gph1
Cip1
Bdh2
Spi1
Ego4
Library
Context
Aim18
Gpx1
Gsy2
ND
Stp4

B

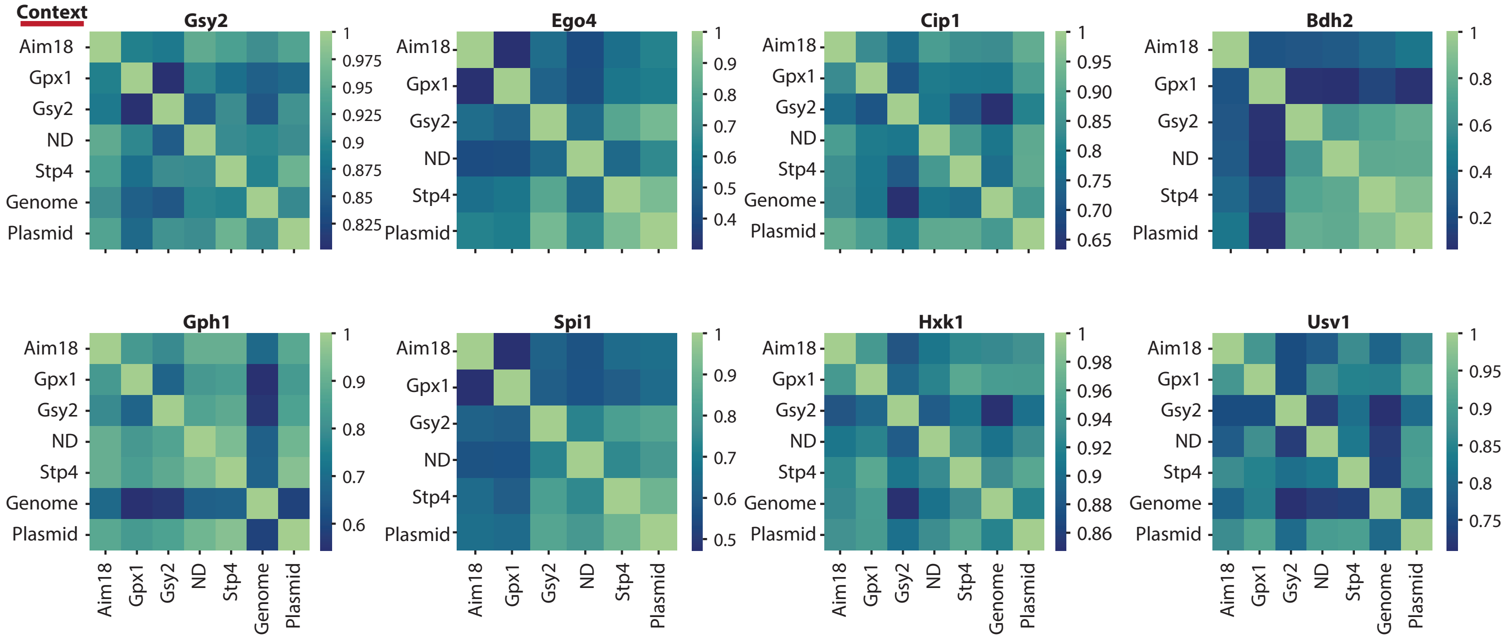
Context
Gsy2
Ego4
Cip1
Bdh2
Gph1
Spi1
Hxk1
Usv1
Aim18
Gpx1
Gsy2
ND
Stp4
Genome
Plasmid

**Figure S6. Changing the surrounding library context affects Msn2 occupancy.**

(A-B) Shown in (A) for each library in each of the tested contexts is the average Msn2 occupancy at the wild-type sequence. Error bars represent the standard error of the mean (SEM) between repeats. (B) Shows the correlation of sequence occupancy between the different contexts for the indicated libraries.

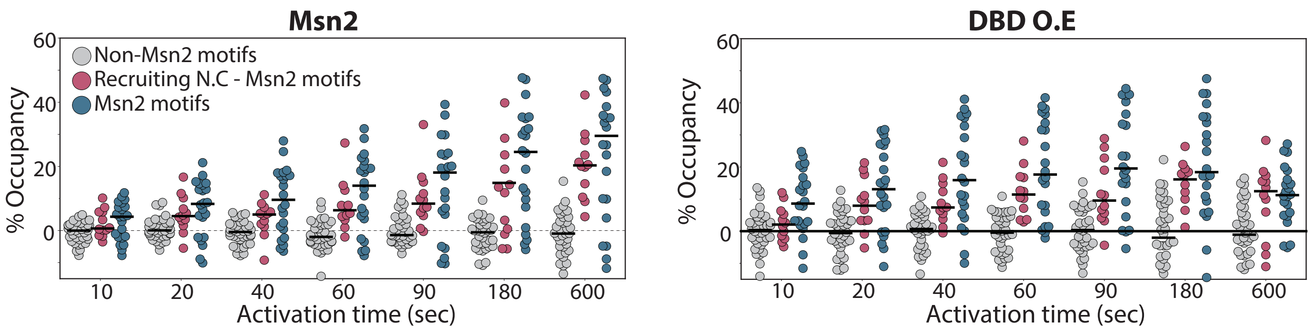
Msn2
Non-Msn2 motifs
Recruiting N.C - Msn2 motifs
Msn2 motifs
% Occupancy
60
40
20
0
10
20
40
60
90
180
600
Activation time (sec)
DBD O.E
% Occupancy
60
40
20
0
10
20
40
60
90
180
600
Activation time (sec)

**Figure S7. Time course showing high correspondence of non-canonical motif preference between Msn2 and an over-expressed mutant containing only its DBD.**

Shown are the occupancies (across all library sequences) of the wild-type Msn2 and of its over-expressed DBD-only mutant as a function of activation times. Sequences are classified based on the presence of a canonical Msn2 motif (AGGGG, blue), a non-canonical motif that contributes to the binding of Msn2 (Hamming distance > 1, pink; as defined in a prior experiment shown in Figure 4, based on the wild-type Msn2 at occupancy at 180 seconds of activation), and non-canonical motifs showing no contribution to the occupancy of Msn2 (gray).

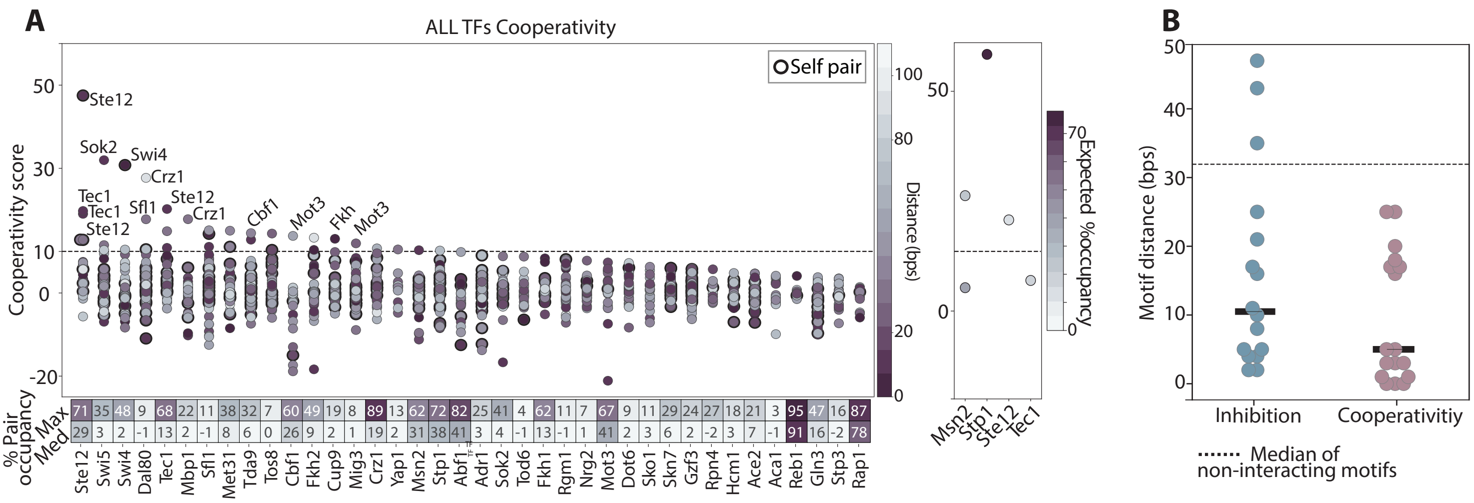
A
ALL TFs Cooperativity
Self pair
Cooperativity score
50
30
10
0
-20
Ste12
Sok2
Swi4
Crz1
Tec1
Tec1
Sfl1
Ste12
Crz1
Ste12
Cbf1
Mot3
Fkh
Mot3
Distance (bps)
100
80
20
0
Expected %occupancy
70
0
% Pair occupancy
Max
Med
71 35 48 9 68 22 11 38 32 7 60 49 19 8 89 13 62 72 82 25 41 4 62 11 7 67 9 11 29 24 27 18 21 3 95 47 16 87
29 3 2 -1 13 2 -1 8 6 0 26 9 2 1 19 2 31 38 41 3 4 -1 13 -1 1 41 2 3 6 2 -2 3 7 -1 91 16 -2 78
Ste12 Swi5 Swi4 Dal80 Tec1 Mbp1 Sfl1 Met31 Tda9 Tos8 Cbf1 Fkh2 Cup9 Mig3 Crz1 Yap1 Msn2 Stp1 Abf1 Adr1 Sok2 Tod6 Fkh1 Rgm1 Nrg2 Mot3 Dot6 Sko1 Skn7 Gzf3 Rpn4 Hcm1 Ace2 Aca1 Reb1 Gln3 Stp3 Rap1
Msn2 Stp1 Ste12 Tec1
B
Motif distance (bps)
50
40
30
20
10
0
Inhibition
Cooperativitiy
Median of non-interacting motifs

**Figure S8. Motif-dependent cooperativity is rare and occurs mainly at proximal motifs.**

(A) *Cooperation among motif pairs is limited:* Shown are the cooperation scores amongst all 1917 motif pairs containing at least one canonical motif of the TFs measured in our screen. Pairs are grouped by the tested TFs, with dots circled in black indicating a pair of motifs belonging to the tested TF. The maximal and median pair occupancies measured for each TF are shown at the bottom. Controls including two variable positions found within the same motif are shown on the right panel with color representing expected pair occupancy. Note the small fraction of cooperating motif pairs.

(B) *The majority of detected interactions are of highly proximal motifs:* Shown are the distances for all motif pairs showing inhibition (left, score<-10) and cooperation (right, score>10) for pairs showing >10% contribution to the TF occupancy. The median values for each group are shown as black lines. The dashed line indicates the median distance between all bound motif pairs showing no interaction.
